# A histone deacetylase network regulates epigenetic reprogramming and viral silencing in HIV infected cells

**DOI:** 10.1101/2022.05.09.491199

**Authors:** Jackson J Peterson, Catherine A Lewis, Samuel D Burgos, Ashokkumar Manickam, Yinyan Xu, Allison A Rowley, Genevieve Clutton, Brian Richardson, Fei Zou, Jeremy M Simon, David M Margolis, Nilu Goonetilleke, Edward P Browne

## Abstract

Approximately 70% of the HIV-1 latent reservoir originates from infections of CD4 T cells that occur in the months near the time of ART initiation, raising the possibility that interventions during this period might prevent reservoir seeding and reduce reservoir size. We identify class 1 histone deacetylase inhibitors (HDACi) as potent agents of latency prevention. Inhibiting HDACs in productively infected cells caused extended maintenance of HIV expression and this activity was associated with persistently elevated H3K9 acetylation and reduced H3K9 methylation at the viral LTR promoter region. HDAC inhibition in HIV-infected CD4 T cells during effector-to-memory transition led to striking changes in the memory phenotype of infected cells. Proviral silencing is accomplished through distinct activities of HDAC1/2 and HDAC3. Thus HDACs regulate a critical gateway process for HIV latency establishment and are required for the development of CD4 T-cell memory subsets that preferentially harbor long-lived, latent provirus.

## Introduction

Antiretroviral therapy (ART) produces rapid and sustained decreases in HIV viremia, yet therapy interruption leads to rapid viral rebound due to a persistent viral reservoir. A key mechanism of HIV persistence is the ability of HIV to enter a state of virological latency in which viral gene expression is silenced, allowing the provirus to remain invisible to the immune system(Chun et al., 1997, 1995; Cohn et al., 2020; Margolis et al., 2020; Perelson et al., 1997). Latency within long-lived memory CD4 T cells permits extended viral persistence, and sporadic reactivation of these cells is sufficient to reseed viremia during treatment interruption(Chun et al., 1997). Longitudinal analysis of the latent reservoir indicates that the reservoir is steadily depleted in ART-treated patients, but this decay rate is too slow for viral clearance within a human lifespan(Chun et al., 1995; Finzi et al., 1999; Perelson et al., 1997). The decay of infected cells is also counterbalanced by the clonal expansion through antigen-driven stimulation and/or homeostatic proliferation(Bailey et al., 2006; Maldarelli et al., 2014; McManus et al., 2019; Wagner et al., 2013). The latent reservoir is thus the main barrier to eradication of HIV in infected individuals. Efforts to eliminate this reservoir have focused on the use of small molecules to reactivate viral gene expression (latency reversing agents, LRAs) in patients durably suppressed by ART, with the hope of rendering infected cells vulnerable to viral cytopathic effect and/or immune clearance. While this approach has achieved some success at inducing viral gene expression in vitro and in vivo, latency reversal methods typically reactivate only a fraction of latent proviruses(Ho et al., 2013; Siliciano and Siliciano, 2021) and have thus-far proven insufficient to measurably reduce reservoir size (Fidler et al., 2020; Gay et al., 2022, 2020; Ho et al., 2013; Kroon et al., 2020; Rasmussen et al., 2014).

The limited efficacy of LRAs is likely due to the multiple layers of transcriptional and epigenetic repression that are added to latent proviruses in a stepwise fashion(Pearson et al., 2008; Rafati et al., 2011). In light of this, we considered an alternative approach: using chemical agents to prevent actively infected cells from entering latency. Such an approach has historically been considered clinically impractical because of observations suggesting that the reservoir is seeded very early during acute infection. However, it was recently shown by longitudinal analysis of HIV sequences during chronic infection that, during viremia, prior to ART initiation, the latent reservoir is more dynamic than previously thought. The majority (∼70%) of both the proviral DNA and replication-competent reservoir is derived from viruses circulating within the last year before ART initiation(Abrahams et al., 2019; Brodin et al., 2006.; Jones et al., 2018). Therefore, ART initiation likely triggers changes in viral and/or immune dynamics in infected individuals, leading to the formation or stabilization of a pool of latently infected cells(Goonetilleke et al., 2019; Margolis et al., 2020). After therapy is initiated, cells harboring intact proviruses are progressively eliminated, likely in large part through low levels of viral gene expression and CD8 T-cell clearance(Einkauf et al., 2022; Falcinelli et al., 2021; Pinzone et al., 2019; White et al., 2022). This model raises the possibility of targeting HIV latency at or before the time of ART initiation to limit the size of the reservoir. For this approach to be successful, detailed information will be required about the molecular mechanisms involved in HIV latency *initiation* rather than *maintenance*.

Activated CD4 T cells are the primary target of HIV-1 infection because they express the CCR5 co-receptor(Shan et al., 2017), undergo effective integration(Zack et al., 1990), and express high levels of transcription factors such as NF-kB that promote viral gene expression(Nabel and Baltimore, 1987). HIV silencing in CD4 T cells is associated with a transition from an activated to resting state (effector-to-memory transition) and correlates with a reduction in host factors that promote HIV gene expression, including NF-kB(Nabel and Baltimore, 1987), the cyclinT1 component of PTEF-b(Budhiraja et al., 2013), and total RNA polymerase II(Shan et al., 2017). Additionally, negative regulators of gene expression are associated with proviral silencing, and repressive modifications to provirus-associated histones such as H3K9me3 and H3K27me3 accumulate on transcriptionally inactive proviruses. Furthermore, intact proviruses that remain after years of therapy tend to be integrated into genomic regions generally enriched in repressive chromatin marks(Battivelli et al., 2018; Einkauf et al., 2022; Jiang et al., 2020). Histone modifications are mediated by a set of host cell enzyme complexes, including histone acetyl transferases (HATs), histone deacetylases (HDACs), histone methyl transferases (HMTs) and lysine demethylases (KDMs)(Boehm and Ott, 2017). These enzymes also play a crucial role in cellular development and fate decisions by generating heritable patterns of gene activation and transcriptional repression that determine cellular phenotype. Importantly, some of these marks can propagate a repressed transcriptional state through cell division, providing a mechanism for epigenetic inheritance of latency.

We recently reported that latently infected cells generated in an *ex vivo* model system of HIV latency exhibit a stable and heritable but reversible state of viral silencing(Jefferys et al., 2021). We therefore used this system to investigate the role of histone-modifying enzyme complexes in the initial establishment of viral latency in CD4 T cells. Our findings indicate that HDACs play a crucial role in the transition of integrated proviruses to a latent state and that Class I HDAC inhibitors (HDACis) potently trap infected cells in a state of continuous viral gene expression. Furthermore, we establish that this activity is associated with epigenetic reprogramming of both the host cell and the provirus through persistent H3K9 acetylation, reduced H3K9 methylation, and inhibition of cellular development into a long-lived stem-cell-like phenotype. We target individual HDACs genetically and pharmacologically to identify distinct contributions from HDAC1/2 and HDAC3 and determine that, while HDAC1/2 are dispensable for the maintenance of proviral latency, these enzymes are required for latency initiation. These data provide important new insights regarding mechanisms of HIV silencing in CD4 T cells and highlight the critical role of HDACs in promoting HIV latency establishment through epigenetic reprogramming. Targeting individual HDACs during ART initiation may block important mechanisms of HIV silencing and limit the formation of a long-lived viral reservoir.

## Results

### HDACis block the establishment of HIV latency in CD4 T cells

To study the initiation of latency in CD4 T cells, we modified an existing reporter virus-based primary CD4 T-cell model of HIV latency(Bradley et al., 2018; Jefferys et al., 2021). The HIV clone (HIV-dreGFP/Thy1.2) used for this model contains a short half-life GFP (dreGFP) within the Env gene and was further modified by insertion of an Internal Ribosome Entry Site (IRES) followed by a murine Thy1.2 gene. (**Figure S1A**). To examine HIV latency initiation, CD4 T cells from healthy donors were activated through the TCR and infected with HIV-dreGFP/Thy1.2 for 48h before magnetic isolation of infected (Thy1.2+) cells (**Figure 1A, S1B**). Isolated Thy1.2+ cells were then expanded 21-28 days in the presence of IL-2 and IL-7. Over this period, we observed a gradual reduction in viral gene expression, with most cells exhibiting undetectable GFP signal by 21d post infection (dpi, **Figure 1B**). Thy1.2 protein levels persisted in many cells that had lost GFP signal, and we hypothesized that slower Thy1.2 protein degradation could allow us to detect infected cells that had recently silenced viral gene expression. We therefore measured the relative decay rates of GFP and Thy1.2 protein in infected cells after halting proviral transcription by CRISPR/Cas9 knockout of the viral Tat gene, and observed that the GFP signal faded with a calculated half-life of 3.3 days, whereas Thy1.2 protein persisted, with a half-life of 10.3 days (**Figure S1C, S1D**). The sustained detection of Thy1.2 protein thus allows us to identify infected cells that are in an early state of latency (Thy1.2+/GFP-). Based on the divergence of stability in these viral reporter proteins we reasoned that the appearance of Thy1.2+/GFP-cells could function as a marker of the emergence of a latently infected population. Furthermore, the frequency of GFP+ cells but not the frequency of Thy1.2 levels responded robustly to LRA treatment (Data not shown). We examined the rate of depletion of Thy1.2+/GFP+ cells with a mixed effects model and observed that the GFP signal was lost from, on average, 2.6% Thy1.2+ cells per day (see statistical methods).

**Figure 1.**
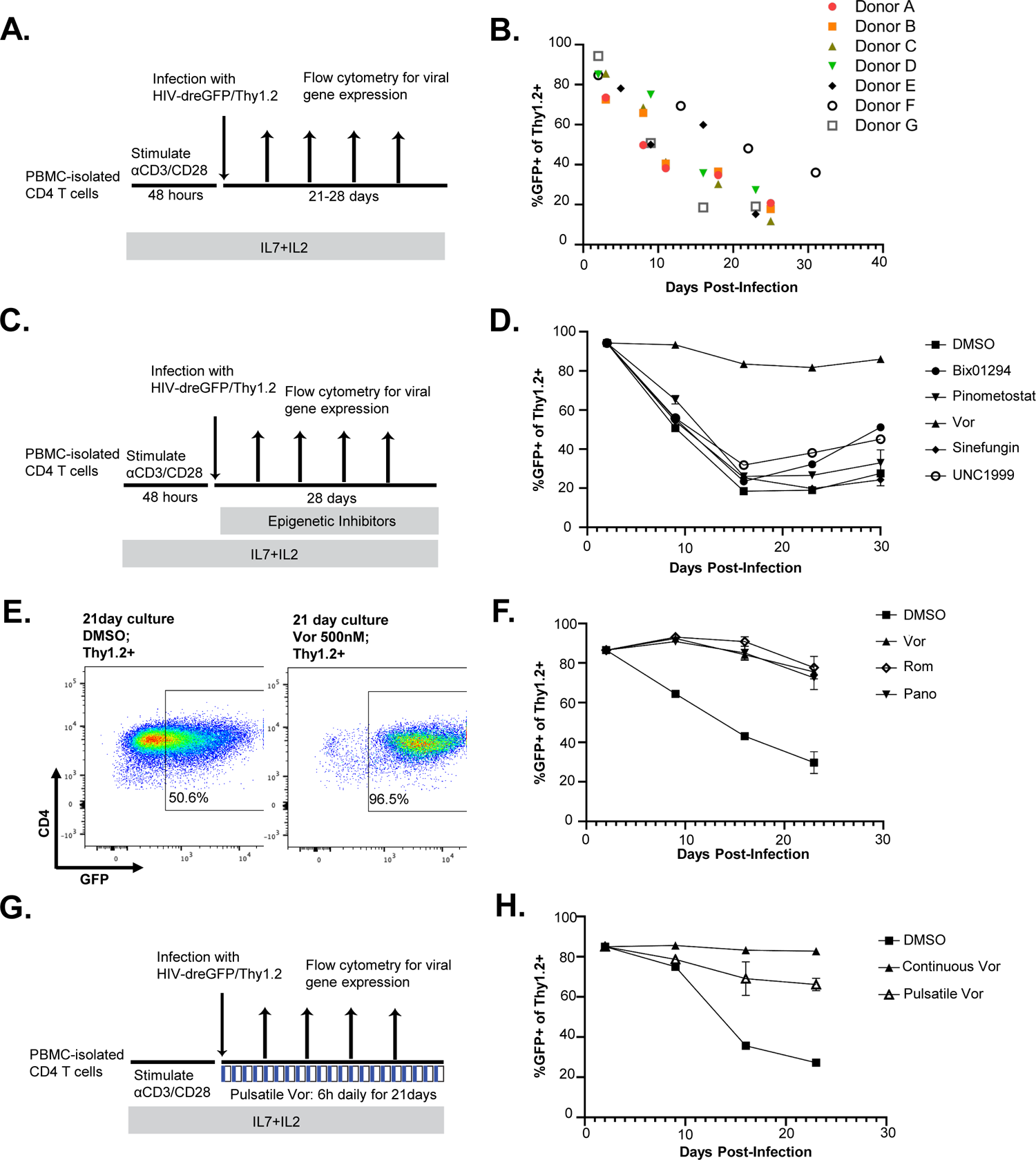
HDACis block HIV latency establishment in CD4 T cells. **A.** Experimental strategy for latency initiation model. Primary CD4 T cells were isolated from PBMCs using negative selection, then activated with beads coupled to anti-CD3 and anti-CD28 antibodies. After 2d, activation, beads were removed, and cells were infected with HIV-dreGFP/Thy1.2. **B**. Progressive decline in viral gene expression as activated cells return to a resting state. The percent of cells productively infected at the indicated times was measured by flow cytometry with gating based on uninfected control cells. Symbols indicate seven individual donors; data here are derived from four independent experiments. **C.** Schematic of epigenetic inhibitor experimental approach. Cells were activated and infected as in Figure 1A and then treated with indicated inhibitors with media refreshed 3 times weekly. **D.** Viral gene expression was measured at different timepoints in cells treated with the indicated inhibitors at the following concentrations: sinefungin 10μM (pan-Methyltransferase inhibitor), vorinostat 500nM (Class I HDACi) BIX01294 250nM (G9a/H3K9me inhibitor), UNC1999 500nM (PRC2/H3K27me inhibitor), pinometostat 10μM (DOT1L/H3K79me inhibitor). **E.** Representative flow cytometry plot of GFP expression in infected cells 14dpi, treated with DMSO vehicle or 500nM vorinostat. **F.** Activated, HIV-dreGFP/Thy1.2-infected CD4 T cells were cultured in the presence of the indicated HDACis as above and percent of productively infected cells was measured weekly by flow cytometry. Vor: Vorinostat, Pan: panobinostat, Rom: Romidepsin. **G.** Schematic of pulsatile vorinostat experiment. Cells were treated continuously or with 500nM vorinostat for 6h daily, with intervening 18h wash-out periods. **H.** Summary of flow cytometry data of viral gene expression over time for continuous or pulsatile vorinostat dosing. Error bars represent standard deviations of two to three technical replicates for all experiments.

We next asked what host cell factors could be responsible for the loss of viral gene expression over time. In particular, we hypothesized that histone modifying enzyme complexes contribute to downregulation of viral gene expression and to the establishment of an epigenetically maintained latent state. To test the role of specific histone-modifying enzymes in HIV silencing in this system, we used a panel of inhibitors of specific enzymes. This panel included sinefungin (pan-HMT inhibitor), UNC1999 (an EZH2/H3K27 methyltransferase inhibitor), BIX01294 (G9a/H3K9 methyltransferase inhibitor), pinometostat (DOT1L/H3K79 methyltransferase inhibitor), and vorinostat (Class I HDACi(Park and Kim, 2020)). First, we treated 2D10 Jurkat cells(Pearson et al., 2008) for 72 hours with efficacious inhibitory doses based on the literature and observed changes to histone methylation and acetylation levels consistent with expected changes (**Figure S1E)**. We then infected activated CD4 T cells with HIV-dreGFP/Thy1.2 and cultured the infected cells in the presence of the inhibitor panel for four weeks (**Figure 1C**). The depletion of the productively infected (Thy1.2+/GFP+) population was measured weekly in primary T cell cultures by flow cytometry (**Figure 1D**). As expected, DMSO (drug solvent) treated cells exhibited progressive down-regulation of HIV gene expression, with approximately 20-50% of cells remaining Thy1.2+/GFP+ at 21 days after infection. Methyltransferase inhibitors UNC1999, Bix01294, pinometostat, and sinefungin had no effect on the loss of GFP or entry of cells into latency. By contrast, vorinostat treatment potently inhibited the development of a latent (Thy1.2+/GFP-) population and maintained the cells in an actively infected GFP+ state (**Figure 1E**). To further investigate this observation, we also examined two other Class I HDACis, panobinostat and romidepsin, and observed that they also prevented the formation of latently infected cells, indicating that this activity is a general feature of class 1 HDACis (**Figure 1F**).

Vorinostat is rapidly cleared during clinical dosing(Ramalingam et al., 2007). We therefore also tested if simulating *in vivo* pharmacokinetics via daily transient vorinostat exposure could prevent HIV latency. Removal of vorinostat containing media for an 18 hour period was previously demonstrated to facilitate clearance of the inhibitor from treated T cells(Clutton et al., 2016). Infected CD4 T cells were exposed to either vehicle only, continuous vorinostat (500nM), or a daily 6h pulse of vorinostat (500nM) followed by an 18h “wash-out” period (**Figure 1G**). Infected cells were monitored for viral gene expression by flow cytometry for 21dpi (**Figure 1H**). As observed above, continuous vorinostat potently prevented the emergence of latently infected cells (3-fold increase in Thy1.2+/GFP+ cells). Pulsatile vorinostat also blocked latency but did not reach the level of efficacy as continuous exposure (2.4-fold increase in Thy1.2+/GFP+ cells).

We also examined whether vorinostat could prevent re-establishment of HIV latency after viral reactivation. We have previously reported that *ex vivo*-generated latently infected CD4 T cells can be reactivated via their TCR to re-express HIV genes but then rapidly re-establish HIV silencing after stimulus removal(Jefferys et al., 2021). We reactivated latently infected cells isolated previously via TCR stimulation and, as expected, stimulated cells rapidly re-expressed GFP, indicating latency reversal. After two days of activation, the TCR stimulus was removed, and cells were exposed to vorinostat (500nM) or vehicle control (**Figure S2A**). Consistent with our previous report, reactivated cells in the vehicle-treated group rapidly re-established latency, with viral gene expression declining 2-fold by 7 days post-activation. By contrast, reactivated cells cultured with vorinostat maintained active viral gene expression (GFP+). Thus, class I HDAC inhibition can prevent both establishment and re-establishment of HIV latency after TCR activation. Overall, we conclude that HDAC activity plays a crucial role in the early stages of latency establishment and that class 1 HDACis block the development of latently infected CD4 T cells from a pool of productively infected cells.

### HIV-infected CD4 T cells maintain viral gene expression after HDACi withdrawal

To further examine the impact of vorinostat on HIV gene expression, we exposed activated HIV-infected cells to a range of vorinostat concentrations for 14 days, followed by an additional 14-day culture in the absence of vorinostat. We observed a dose-dependent relationship between vorinostat concentration and latency prevention, with a consistent trend toward latency reduction observed even in the lowest dose of vorinostat (125nM) (**Figure S2B**). Based on the measurable latency prevention observed at this low dose of vorinostat, we sought to compare the activity of HDACis in latency prevention versus latency reversal. We tested HDACis for latency reversing activity in two latently infected Jurkat cell lines - one with a provirus that is highly responsive to LRAs (2D10 cells(Pearson et al., 2008), **Figure S2C**) and another that is relatively unresponsive to LRAs (N6 cells(Nixon et al., 2020), **Figure S2D**). 2D10 cells were responsive to HDACis but only at relatively high doses (vorinostat EC50: 671nM), while N6 cells were almost completely unresponsive to the HDACis in the range tested. These data are comparable to a previous analysis across diverse latent cell models, where maximal latency reversal was observed at 1μM vorinostat(Spina et al., 2013). We examined whether latency prevention in primary T cells was durable after withdrawal of the inhibitor and observed that vorinostat-treated cells maintained significantly elevated HIV expression, even 14d post-vorinostat removal (**Figure S2E**). Thus, a transient window of vorinostat exposure by infected cells during the effector-to-memory transition can lead to protracted viral gene expression post-exposure, even for a low concentration (125nM). Thus, we speculate that latency prevention in primary cells is distinct from latency reversal and is characterized by a higher level of vulnerability to HDACis. Additionally, we propose that transient exposure of actively infected CD4 T cells to HDACis during transition from an activated to a resting state can have a long-lasting impact on viral gene expression, possibly through interfering with the establishment of a repressive epigenetic state at the provirus.

### T cell activation elevates global histone acetylation and HDACis partially maintain this state

To investigate mechanisms of prolonged viral gene expression in the presence of vorinostat, we first examined the levels of acetyl-histone modifications in CD4 T cells after HDACi treatment or after T-cell activation. Using techniques to fractionate primary CD4 T cells into cytoplasmic, soluble nuclear, and chromatin-bound extracts (**Figure S2D**) we determined that the levels of acetyl-Histone H3 Lysine 9 (H3K9ac) and acetyl-Histone H3 Lysine 27 (H3K27ac) were highly elevated after 72 hours of either vorinostat (**Figure 2A**) or T-cell activation (anti-CD3/CD28 stimulation) (**Figure 2B**). We speculated that T-cell activation increases global histone acetylation through histone acetyl transferase (HAT, ie: P300, CBP) recruitment to TCR-responsive promoters and enhancers(Yukawa et al., 2019), and hypothesized that HDACi treatment may maintain elevated global chromatin acetylation levels after T cell activation. To examine this hypothesis, CD4 T cells were activated for 72 hours, allowed to grow for an additional 72 hours, and subsequently treated with vorinostat for 14 days or control vehicle (DMSO). In contrast to the robust effect we observed with vorinostat treatment or TCR activation, longer-term (7 or 14 days) vorinostat treatment of activated T cells led to relatively modest changes in total levels of H3K9ac and H3K27ac (**Figure 2C**). Specifically, global H3K27Ac levels were not affected by 14d vorinostat, while H3K9Ac were only modestly increased. Thus exposure of CD4 T cells to vorinostat during a return to rest maintains viral gene expression independently of large global changes in histone acetylation.

**Figure 2:**
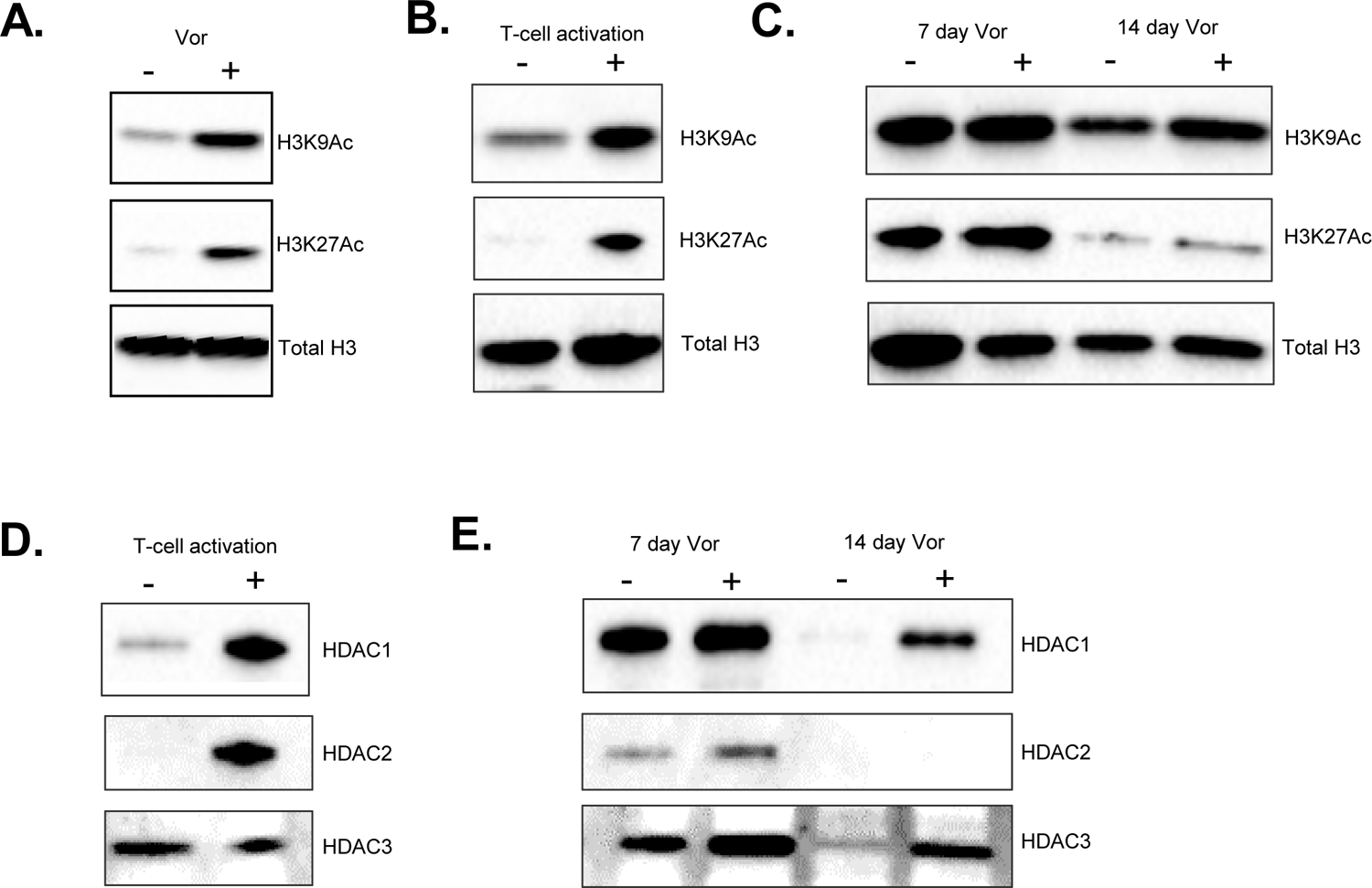
HDACi exposure during return of activated CD4 T cells to rest leads to persistently elevated global H3K9 acetylation. **A-E.** Immunoblots of chromatin-associated protein from HIV-dreGFP/Thy1.2 infected cells under the indicated conditions. Cells were cultured in the presence of 100U/mL IL-2 and 2.5ng/mL IL-7. Chromatin fractions are proteins liberated by DNase treatment after elution of cytoplasmic and soluble nuclear pellets. **A**. Immunoblots of chromatin fractions after 72 hours of DMSO (“-”,1:1000) or vorinostat (“+”, 500nM) treatment. **B**,**D**. Immunoblots of chromatin fractions after 72 hours of culture (“-“) or T cell activation (“+”,anti-CD3/CD28 dynabeads). **C,E.** Immunoblots of chromatin fractions after activated T cells were treated for 7 or 14 days with (“+”, 500nM) or without (“-“, 1:1000 DMSO) vorinostat. Cell media was replaced three times weekly with fresh media containing cytokines and DMSO or vorinostat.

HDAC activity is regulated by the recruitment of HDACs to DNA by specific transcription factors, and thus even though Class I HDAC expression levels hev been previously reported to be generally unaffected by T-cell activation,(Keedy et al., 2009) we reasoned that T-cell activation may influence the levels of chromatin-associated HDACs. We thus examined the levels of Class I HDACs in chromatin extracts from CD4 T cells before and after T-cell stimulation. Whereas a baseline level of HDAC3 was constitutively associated with chromatin with or without T-cell activation, chromatin-associated HDAC1 and HDAC2 were nearly absent from the chromatin fraction at rest, but were robustly recruited by T-cell activation (**Figure 2D**). As the cells returned to rest over 14 days, chromatin-bound HDAC1 and HDAC2 levels exhibited a progressive decline indicating a restoration of the low HDAC1/2 homeostasis, whereas chromatin-bound HDAC3 levels were relatively consistent. Notably, long-term vorinostat treatment was associated with increased levels of chromatin-associated HDAC1 (**Figure 2E**). Overall, these data indicate that chromatin bound levels of HDAC1 and HDAC2 are highly dynamic soon after T cell activation, exhibiting a rapid and dramatic increase followed by a gradual depletion and return to baseline levels.

### Persistently elevated H3K9 acetylation blocks proviral LTR methylation

We next sought to clarify the impact of latency prevention through Class I HDAC inhibition on the abundance of activating and repressive histone marks on the HIV provirus. We hypothesized that, although continuous vorinostat exposure does not lead to robust global changes in histone acetylation and methylation, it could promote more pronounced local changes within the HIV provirus. To measure genome-wide and provirus-associated histone modifications in the limited cell numbers achievable in primary T cell cultures, we used Cleavage Under Target and Release Under Nuclease (CUT&RUN)(Meers et al., 2019; Skene and Henikoff, 2017) to profile histone modifications associated with active (H3K4me3, H3K27ac, and H3K9ac) and repressed (H3K9me3 and H3K27me3) regions of the genome in infected cells that had been cultured with or without vorinostat for 21 days post activation (**Figure 3A, 3B**). We also examined genome-wide binding patterns of total RNA Polymerase II (RPD1 subunit, RNApol2) and RNA Polymerase II phosphorylated on Ser 2 of the heptameric repeat region (RNApol2 P-S2), a modification associated with polymerase processivity.

**Figure 3:**
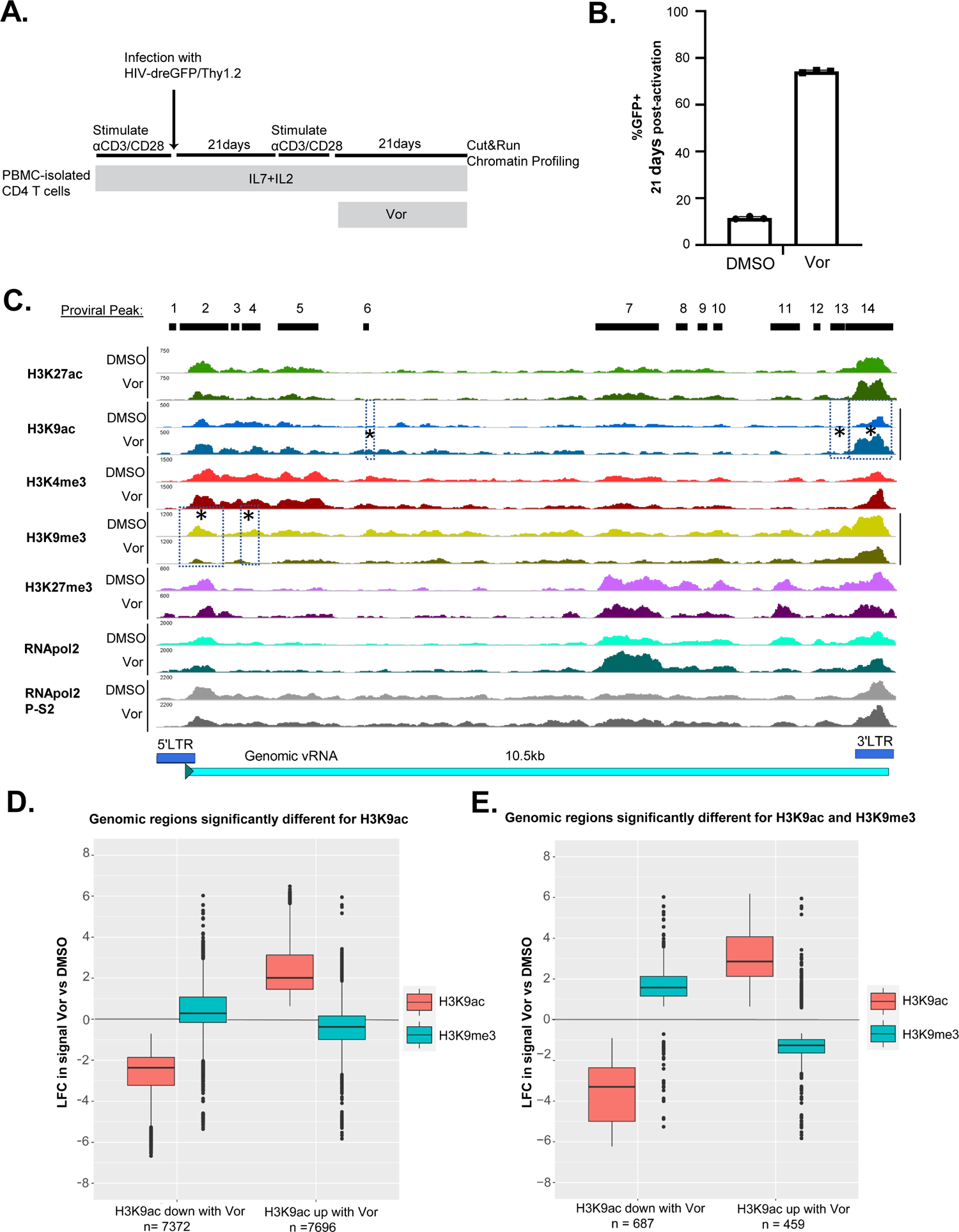
Persistently elevated H3K9 acetylation blocks methylation of the proviral LTR. **A.** Experimental overview. CD4 T cells were activated and infected with HIV-dreGFP/Thy1.2, Infected (Thy1.2+) cells were isolated, and cells were cultured for 21d. Cells were re-stimulated with anti-CD3/anti-CD28 beads for 2d and cultured in the presence of 330nM vorinostat for 21d before CUT&RUN chromatin profiling against the indicated targets. **B.** Bar graph of frequency of GFP+ cells measured by flow cytometry after 3 weeks of culture in vorinostat. **C.** Normalized read density aggregated (see methods) from 2-3 CUT&RUN technical replicates for each condition were plotted across the HIV proviral genome. Top black bars: Union set of regions identified across antibody and treatment conditions. Bottom: HIV proviral schematic. Regions found to exhibit a significant (p<0.01) difference in enrichment between vehicle and vorinostat treatments are marked with an asterisk. **D,E** Box and whisker plots of H3K9ac and H3K9me3 signal at genomic sites that exhibited a significant difference in enrichment between vehicle and vorinostat treatments for H3K9ac only; (**D**) or a subset containing regions significantly changed for both H3K9ac and H3K9me3 (**E**). Positively (left) and negatively (right) regulated H3K9ac peaks are plotted, and relative fold-change of H3K9me3 signal is compared at these same sites. Box indicates interquartile range and line indicates median fold-change of peaks; “LFC” indicates log_10_ fold-change between treatment conditions. Statistical analyses were performed using csaw and EdgeR with a significance threshold of p<0.01 using a Negative Binomial Generalized Linear model, see methods.

Alignment of the data to a combined human and HIV proviral genome indicated clear enrichment of each mark at specific peaks. We detected a total of 1,460,525 peaks across both the host and proviral genomes that were enriched in at least one of the antibody and treatment conditions. We examined the correlation of signal at these sites between each CUT&RUN condition and found that replicate samples clustered by antibody target, and that activating factors (H3K4me3, H3K9Ac, H3K27Ac) generally correlated with each other, whereas H3K9me3 and H3K27me3 (associated with repressive chromatin) clustered separately (**Figure S3**). Samples treated with vorinostat clustered together in some but not all antibody conditions. Looking specifically at the HIV provirus, we observed 14 distinct enriched regions (peaks numbered 1-14) across all conditions/antibodies (**Figure 3C**). Highlighting the importance of the 5’ and 3’ LTR regions for epigenetic regulation, peaks 1 and 2 overlapped with the 5’ LTR, while peak 14 covered the 3’ LTR. The primary T cell model employed in this experiment contains a diverse set of integration sites and thus the measurements by CUT&RUN should be interpreted as a bulk measurement of the overall ensemble of proviruses in the sample. Intriguingly regulatory marks were at least as abundant at the 3’ LTR of the virus as at the 5’ LTR. This finding remains unexplained but is consistent with our previous measurements identifying a large DNA accessibility footprint at the 3’ LTR(Jefferys et al., 2021).

To examine the effect of vorinostat on binding of these factors, we identified regions across the host and proviral genomes whose signal significantly (p<0.01, csaw sliding-window method(Lun and Smyth, 2016)) differed between vorinostat and control conditions. Examining marks within the 14 proviral peaks between the vorinostat-treated and DMSO-treated cells, we determined that CUT&RUN signal for most antibodies at most regions were not significantly different between vorinostat treated and untreated conditions. However, peaks 6,13, and 14 exhibited significantly elevated H3K9ac occupancy in the vorinostat-treated cells (**Figure 3C**). We also observed a significant reduction in H3K9me3 associated with peaks 2 and 4 in vorinostat-treated cells. Based on these observations, we conclude that vorinostat exposure during a return to a resting state leads to persistently elevated H3K9 acetylation at multiple proviral sites, particularly in the 3’ LTR, as well as reduced H3K9me3. We hypothesize that H3K9me3 is likely blocked by persistent H3K9Ac levels, since HMTs require unmodified H3K9 as a substrate. According to this model, locking an acetyl group on the proviral H3K9 residues in activated T cells while both histone acetylation and HDAC levels are elevated would prevent the subsequent addition of repressive marks that can mediate stable and heritable epigenetic silencing.

To test whether the reciprocity between H3K9ac and H3K9me3 at the HIV provirus reflected a more general principle of genomic regulation, we further examined the genome-wide distribution of the H3K9ac and H3K9me3 marks after vorinostat treatment. Globally we identified 15,068 genomic sites with significant changes in H3K9ac levels and 84,313 sites with significant changes in H3K9me3. With vorinostat treatment, we identified 15,068 genomic peaks with significant (p<0.05) changes in H3K9ac levels and 84,313 peaks with significant changes in H3K9me3. Across all these dynamic regions, we observed a modest but significant inverse-correlation of these two marks (**Figure 3D, Figure S3**). The inverse correlation between H3K9ac and H3K9me3 was more pronounced when we specifically examined the 1,146 sites that exhibited a significant change following vorinostat treatment in both marks simultaneously (**Figure 3E**). These data support the existence of a reciprocal regulatory switch between these marks; H3K9ac removal by HDACs as activated T cells return to a resting state serves as a ‘gatekeeping’ change that licenses the subsequent deposition of H3K9 methylation at a subset of loci, including the HIV provirus. We speculate that maintenance of a permissive chromatin state by HDACi in combination with the positive-feedback mechanisms of the HIV transactivator protein Tat may together account for the observed persistence of viral gene expression, although notably we failed to see evidence of increased HIV-associated RNApol2 P-S2 which we had anticipated would serve as a marker of Tat function.

### HDAC inhibition during return of activated T cells return to rest causes changes in the host cell transcriptome and emergence of a memory phenotype

Given the impact of vorinostat exposure during effector to memory transition on the global abundance of histone modifications, we next investigated the broader impact of this treatment on the host cell. As cellular transcription factors are frequently regulated by acetylation, we hypothesized that, even in the absence of large global changes in histone acetylation, HDACis might promote an altered cellular transcriptional program that affects viral gene expression. We analyzed the transcriptomes and CD4 T-cell memory phenotype of CD4 T cells that had been activated, infected, then exposed to vorinostat or vehicle control for 14 days (**Figure 4A**). As expected, we observed an increase in viral gene expression measured by both flow cytometry (**Figure 4B**), and by counting reads aligning to the HIV genome (**Figure 4C**). Differential-expression analysis of host-transcripts revealed 1322 downregulated and 1978 upregulated genes detectable with continuous vorinostat exposure (**Figure 4D**). To broadly characterize differentially expressed genes we used Hallmark Pathway gene set enrichment analysis of differentially expressed genes to acquire insight into host cell processes that may underly the observed changes in memory phenotype(Liberzon et al., 2015). This analysis revealed that upregulated genes included those involved in TNFα signaling through NF-κB, Hedgehog signaling, the p53 pathway, the Interferon-γ response, and Apical Junction (**Figure 4E**). In downregulated transcripts (**Figure 4F**), significantly enriched terms were involved in G2M checkpoint regulation, E2F target genes, mitotic spindle formation, hypoxia, MTORC1 signaling, and allograft rejection. The robust enrichment of cell cycle-associated transcripts in downregulated genes is consistent with the reported essential role of HDAC1 and HDAC2 in cell cycle progression(Jamaladdin et al., 2014; Wilting et al., 2010) and the decreased proliferation we observed in vorinostat-treated cultures (Data not shown).

**Figure 4:**
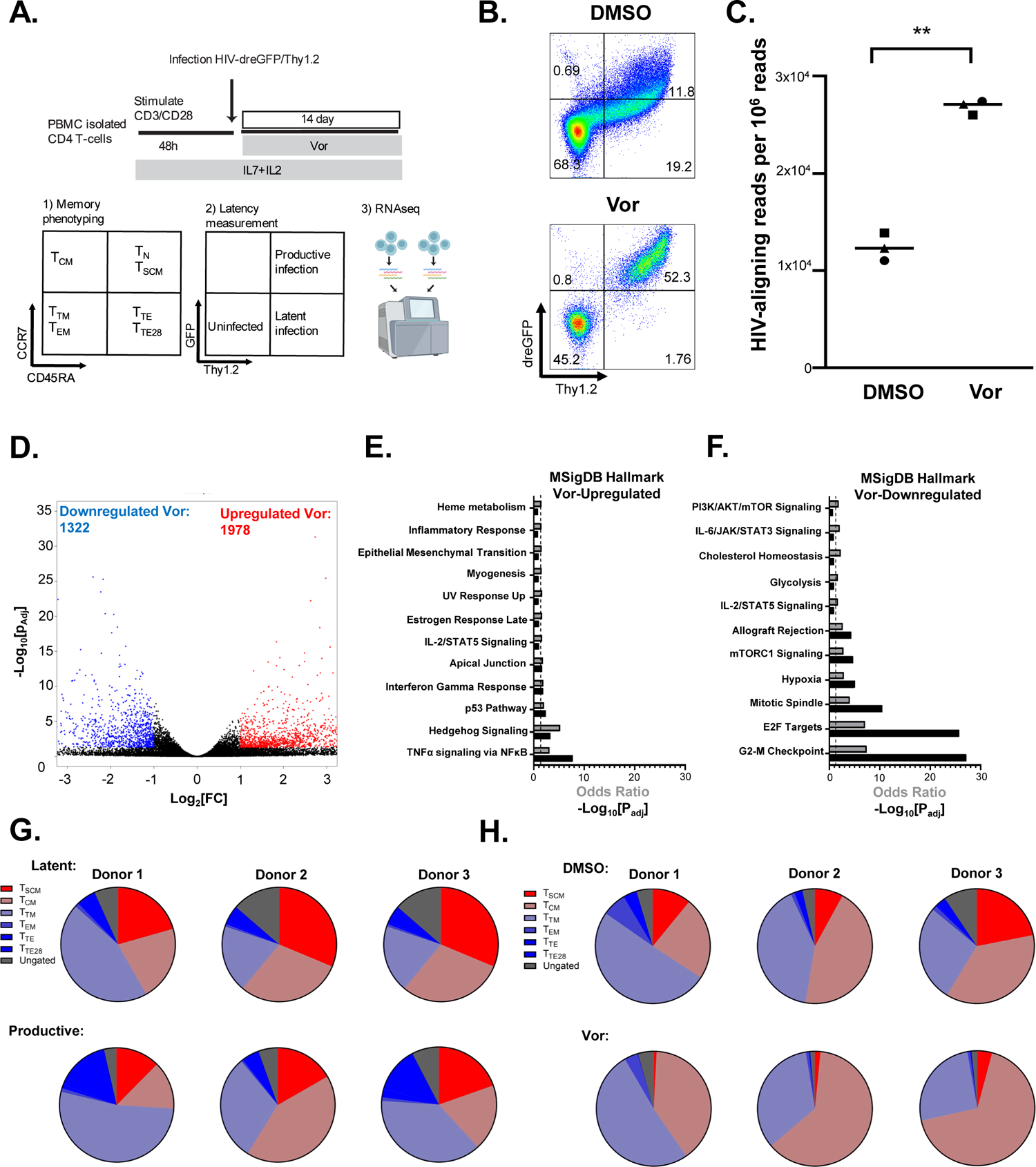
HDAC inhibition during effector-memory transition alters the transcriptome and cell-fate of HIV-infected CD4 T cells. **A.** Schematic of integrated host-cell characterization experiment. CD4 T cells from three independent donors were activated via their TCR and then infected with HIV-dreGFP/Thy1.2 for 48h. Infected cells were then cultured for an additional 14d with vorinostat (500nM) or control vehicle (DMSO 0.1%). Cells were then profiled by RNAseq **(C,D,E,F)** and flow-cytometry (**B,G,H**). **B.** Flow cytometry for viral gene expression measured by Thy1.2 and GFP signal. **C.** Viral gene expression was quantified in each sample by alignment to the HIV-dreGFP/Thy1.2 genome and normalized to total sample read-depth. Statistical comparison with generalized estimating equations, see methods, ***=p<0.0001. **D.** Volcano plot indicating differentially expressed genes in cells exposed to vorinostat as activated T cells return to a resting state. Red: upregulated genes (Padj<0.05, Log_2_ Fold change>1). Blue: downregulated genes (P_adj_<0.05, Log_2_ Fold change<-1). **E,F.** Hallmark analysis of pathways enriched (p<0.05) in upregulated (**E**) and downregulated (**F**) (|Log2FC|>1, P_adj_<0.05) transcripts. **G**. Summary of memory cell subset distribution in cells gated on latent (top pie-charts; Thy1.2+/GFP-) and productive (bottom pie charts; Thy1.2+/GFP+) cell populations. Gating performed as described in (**Figure S4**). **H.** Pie charts as in (**G**) but comparing vehicle (top)- and vorinostat (bottom)-treated cells.

Memory T cells can be differentiated based on distinct immune phenotypic markers defining features such as anatomical trafficking (ie: the lymphatic homing receptor CCR7), whether the cells are antigenically experienced (ie: CD95), or by expression of receptors that grant the capacity to respond to important cytokines (eg IL-7 receptor, CD127). The prevailing system to classify memory T cells relies on surface markers that can be used to distinguish the differentiation state of the T cell as follows: naïve (T_N_) or stem cell-like memory cells (T_SCM_), central memory (T_CM_), T transitional memory (T_TM_), effector memory (T_EM_), and terminal effector (T_TE_) cell subsets. Generally, CD4 T cells further along in this stated order possess more effector functions but lose proliferative and self-renewal potential. These changes in CD4 T cells are guided by transcriptional programs that define and maintain subset identity; importantly, studies show that the latent reservoir is enriched in longer-lived memory cells(Manganaro et al., 2018; Soriano-Sarabia et al., 2014) (T_CM_ and T_SCM_) and that cells with effector character more readily express viral gene products(Bradley et al., 2018; Kulpa et al., 2019; Wonderlich et al., 2019), although this association is not detected in every study of PLWH and may depend on the cohort or method of latent reservoir quantification(Duette et al., 2022; Kwon et al., 2020). We therefore sought to define the association between viral latency and CD4 T-cell memory subsets in our model system. CD4 T cells from healthy donors were activated and infected with HIV-dreGFP/Thy1.2 and cultured for 14 days before being characterized by a flow cytometry panel of T-cell memory markers. By sequentially gating on CCR7 and CD45RA, then CD28 and CD95, we categorized cells into T_N_, T_CM_, T_SCM_, T_TM_, T_EM_ and T_TE_ cells (**Figure S4A-S4D**)(Mahnke et al., 2013). Most cells exhibited T_CM_, T_SCM_, and T_TM_ phenotypes, with minor subpopulations of T_EM_ and T_TE_ cells. Classically, T_TE_ cells are defined as CD28-; however, in our model system, we identified that most CCR7-/CD45RA-cells were CD28+, a population we notate as T_TE28_. By examining the level of viral gene expression in each subset, we observed differences in the distribution of latently infected (Thy1.2+/GFP-) versus productively infected (Thy1.2+/GFP+) cells across subsets. Specifically, latent infection was observed more frequently enriched in cells exhibiting a T_SCM_ or T_CM_ phenotype, while the T_TM_ and T_TE28_ subsets were preferentially enriched in productively infected cells (**Figure 4G**). We also examined the distribution of latency based on expression of the memory marker CD127. Consistent with a recent report(Hsiao et al., 2020), latency was more frequent in CD127+ cells, further suggesting that latency preferentially occurs in cells that adopt a long-lived memory phenotype (**Figure S5A**). We also observed that PD-1, a marker of both T cell exhaustion and of T cell activation, was enriched in productively infected cells (**Figure S5B**).

Interestingly, we noted that vorinostat-treated CD4 T cells exhibited a consistently altered memory phenotype (**Figure 4H**). In particular vorinostat exposure resulted in a reduced fraction of cells with a T_SCM_ phenotype (CCR7+, CD45RA+, CD95+, CD28+) and a shift to a T_CM_ phenotype. This phenomenon was true for all cells, regardless of infection (Thy1.2+) status, indicating that the observed subset reprogramming was not specific to HIV-infected cells. We also examined the frequency of CD127 and PD-1 expression and found no changes in CD127+ cells and a slight reduction in PD1+ cells with vorinostat (**Figure S5C-S5F**). We observed that vorinostat exposure did not affect T cell activation, as measured by PD-1 (**Figure S5F**) or HLA-DR expression (**Figure S5G**). Overall, these data demonstrate that HIV silencing across the CD4 T-cell population was not random but was associated with the identities adopted by the individual cells as they return to rest. In particular, infection of cells that develop into naïve-like, self-renewing cells such as T_SCM_ and T_CM_ subsets with elevated CD127 expression is more likely to result in viral silencing and latency. The preferential silencing of the provirus in long-lived memory-like cells is likely an important mechanism that contributes to viral persistence and the capacity for clonal expansion of infected cells(Spina et al., 1997; Murray et al., 2016; Pinzone et al., 2019; Simonetti et al., 2021). Furthermore, we observe that vorinostat exposure during the effector to memory transition significantly alters the subset identity distribution of the overall population. This identity reprogramming may contribute to the impact of vorinostat on viral gene expression in this system. It should be noted, however, that both latent and productive infection were identified in each CD4 T cell memory subset, demonstrating that the host-cell environment alone is not solely deterministic for viral gene expression. Nevertheless, these observations are consistent with the notion that epigenetic programming during cell fate decisions can regulate HIV entry into latency. The finding that HDAC inhibition as activated CD4 T-cells transition to a resting state leads to dysregulated memory formation suggests that HDACi could disrupt not only viral silencing but also the formation of long-lived memory cells, a process we have previously postulated is a targetable component of the stabilization of the latent reservoir in patients who have initiated ART(Goonetilleke et al., 2019).

### HDAC1/2 and HDAC3 play essential and distinct roles in HIV silencing in CD4 T cells

We next wished to identify the cellular targets by which vorinostat prevents HIV silencing in CD4 T cells during effector to memory transition. Vorinostat inhibits Class I HDACs (HDAC1, HDAC2, HDAC3 and HDAC8). However, the contribution and roles of individual HDACs are largely unknown in primary T cells and have not been defined in the context of latency initiation(Keedy et al., 2009). Because of the high homology of HDAC1 and HDAC2 in the enzyme activation site, selective inhibitors for HDAC1 or HDAC2 have not been developed. Thus, we examined the impact of novel small molecules that inhibit both HDAC1 and HDAC2 (HDAC1/2i) or HDAC3 (HDAC3i) on HIV latency initiation in CD4 T cells(Clausen et al., 2020; J. Liu et al., 2020; Yu et al., 2021). Activated CD4 T cells were infected with HIV-dreGFP/Thy1.2 and cultured for 14d in the presence of each inhibitor, and viral gene expression was measured by flow cytometry (**Figure 5A**). Importantly, we observed dose-responsive latency prevention with the HDAC1/2i with a similar potency to vorinostat, indicating a key role for HDAC1 or HDAC2 (**Figure 5B**). By contrast, an HDAC3i (Merck HDACi compound 38(J. Liu et al., 2020)) prevented HIV silencing to a lower extent (**Figure 5B**). In time course latency initiation experiments we saw that the HDAC1/2i phenocopied vorinostat at preventing latency initiation (**Figure 5C**) an observation that was confirmed with measurements at 21 days post-infection across 5 separate donors (**Figure 5D**) where statistically significant increases in Thy1.2+;GFP+ cells were observed with both vorinostat (Wilcoxon ranked-sum test, p_adj_=0.035) and HDAC1/2i (p_adj_=0.035) but not with HDAC3i (p_adj_=0.278). A structurally distinct HDAC3i (RGFP966(Malvaez et al., 2013)) also failed to maintain viral gene expression in latency re-initiation experiments (**Figure S6A**). Remarkably, for HDAC1/2i-treated cells, we observed a progressive increase in viral gene expression intensity over time, significantly beyond that induced by vorinostat (**Figure 5E**). Indeed, HDAC1/2i-treated cells had an approximately two-fold higher GFP intensity than cells immediately after activation and infection, the typical point of maximal GFP signal. We reason that specific targeting of HDAC1 and HDAC2 may be more effective or tolerable than a broader block to class 1 HDACs with respect to preventing HIV silencing in CD4 T cells.

**Figure 5:**
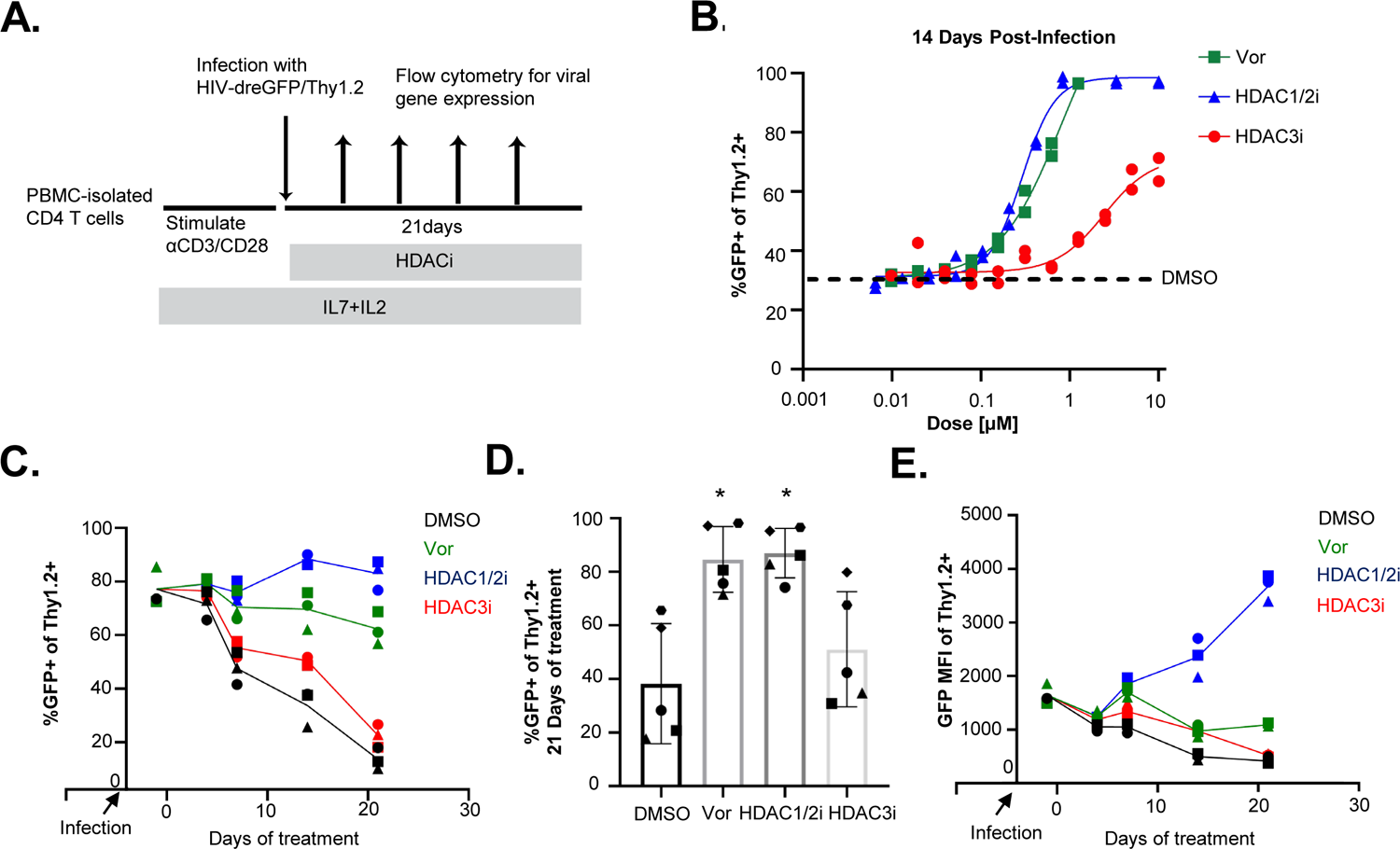
Class I HDAC selective inhibitors reveal essential role of HDAC1 and 2 in latency initiation. **A.** Schematic of HDAC-selective inhibitor experiments. HIV-dreGFP/Thy1.2-infected primary CD4 T cells were cultured in the presence of a dose curve of an HDAC1 and HDAC2 inhibitor (HDAC1/2i), an HDAC3 inhibitor (HDAC3i), vorinostat (Vor), or control vehicle (DMSO 0.1%) for 14dpi, and productive viral gene expression was examined by flow cytometry. **B.** Dose response curve showing proportion of cells with productive viral gene expression 14d post treatment with the indicated inhibitors. Plots are from one representative donor conducted in technical triplicate. **C.** Time course flow cytometry experiment in HIV-dreGFP/Thy1.2-infected primary CD4 T cells cultured with the indicated inhibitors. Vor: vorinostat 500nM, HDAC1/2i: 800nM; HDAC3i:3μM. Points represent three donors assessed in parallel **D**. Bar graph of productive viral gene expression measured after 21 days of treatment with indicated inhibitors; symbols indicate data from five individual CD4 T cell donors and error bars indicate standard deviation. Statistical comparison was performed between DMSO treatment condition and all other treatment conditions with the Wilcoxon ranked-sum test with a Bonferroni adjustment for multiple comparisons; *=p_adj_<0.05. **E**. Mean fluorescent intensity (MFI) of GFP signal in Thy1.2+ cells in data from (**C**).

The prevention of latency initiation observed with HDAC1/2 inhibition came as a surprise due to our previous work that found that targeting HDAC1 and HDAC2 alone or together did not robustly reverse latency(Barton et al., 2014; Keedy et al., 2009). We confirmed these previous observations by testing the HDAC1/2i in latency reversal experiments in both the 2D10 cell-line model of latency and in primary T cells three weeks after infection with HIV-dreGFP/Thy1.2. Unlike in latency prevention experiments, where HDAC1/2i reproduced the effect of pan-Class I HDAC inhibition with vorinostat, HDAC1/2i failed to reverse latency in either of the tested models (**Figure S6B-S6D**). The HDAC3i compound 38 also failed to reverse latency, whereas RGFP966 exhibited activity in 2D10 cells but not in latently infected primary T cells. These results are consistent with the hypothesis that the mechanisms governing latency initiation are distinct from mechanisms that enforce latency maintenance.

We next sought to further explore the roles of individual HDACs in latency initiation by using CRISPR/Cas9 targeting to knockout individual Class I HDACs in HIV-infected CD4 T cells (**Figure 6A**). We optimized conditions for CD4 T-cell nucleofection with sgRNA/Cas9 ribonucleoprotein complexes, enabling robust depletion of HDAC1, HDAC2, HDAC3, HDAC8, or of both HDAC1 and HDAC2 by 7 days after nucleofection, as measured by immunoblot (**Figure 6B**). HDAC2 expression increased after HDAC1 knockout, likely reflecting a regulatory feedback loop (**Figure 6B**). Interestingly, knockout of HDAC3 significantly increased the proportion of actively infected (Thy1.2+/GFP+) cells and HDAC1 knockout trended towards increasing the proportion of Thy1.2+/GFP+ cells, whereas targeting HDAC2 or HDAC8 individually had no impact on viral gene expression (**Figure 6C**). Since HDAC1 and HDAC2 are highly homologous and engaged in the same protein complexes, we hypothesized that HDAC2 upregulation after HDAC1 knockout may be sufficient to compensate for HDAC1 knockout. Thus, we targeted both HDAC1 and HDAC2 with CRISPR/Cas9 in combination and observed a larger effect on latency initiation, albeit not statistically significant due to smaller sample size. Additionally, similar to observations made in genetically engineered mouse model experiments(Wilting et al., 2010) we saw that growth arrest was seen with combined HDAC1 and HDAC2 knockout but not with either knockout alone; this phenomenon likely leads to a loss of double-knockout cells over time.

**Figure 6:**
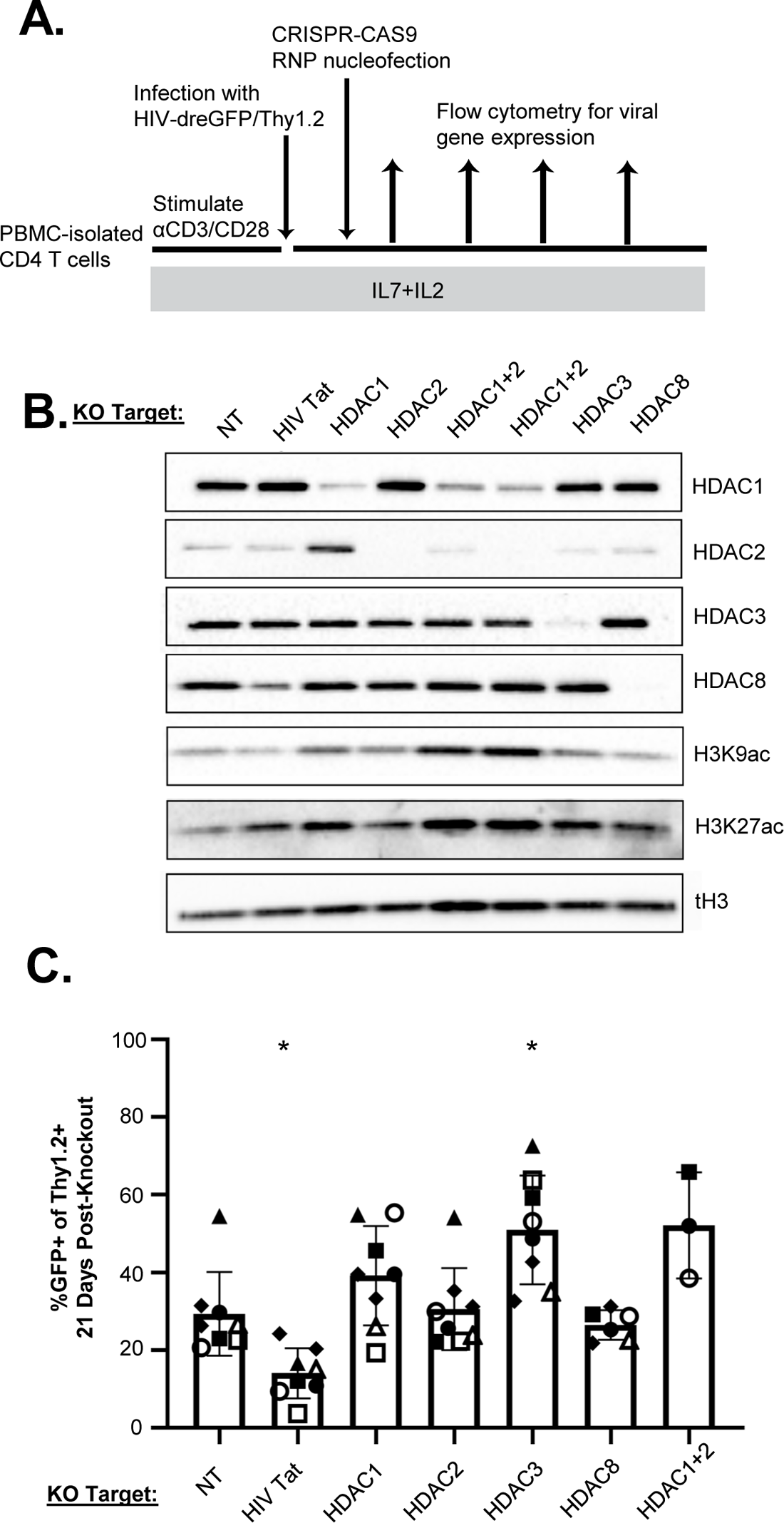
CRISPR/CAS9 knockout of class I HDACs reveals requirement for HDAC3 in latency initiation. **A.** Schematic overview of HDAC CRISPR/Cas9 knockout experimental design. CD4 T cells were activated, infected with HIV-dreGFP/Thy1.2, then nucleofected with Cas9/sgRNA RNPs with either a non-targeting sgRNA or sgRNA directed against HIV-1 Tat, HDAC1, HDAC2, HDAC3, HDAC8, or both HDAC1 and HDAC2. **B.** Immunoblot of whole cell lysates from cells 7d after sgRNA/Cas9 RNP nucleofection. NT indicates non-targeting CRISPR control condition. **C.** Bar graph of productive viral gene expression measured in cells 21d after sgRNA/Cas9 RNP nucleofection. Symbols indicate data from eight individual CD4 T cell donors and error bars indicate standard deviation. Samples were excluded if less than 50% depletion of target was measured by immunoblot; five HDAC8 knockout donors and three HDAC1+2 double knockout donors were assessed. Statistical comparison was performed between the non-targeting CRISPR control and all other treatment conditions with the Wilcoxon ranked-sum test with a Bonferroni adjustment for multiple comparisons; *=p_adj_<0.05.

Taken together, the inhibitor and CRISPR knockout data suggest that HDAC1 and HDAC3 play essential but mechanistically distinct roles in latency initiation in CD4 T cells, and that HDAC2 can partially compensate for HDAC1 loss because of overlapping functional roles. Given the impact of HDAC3 CRISPR/Cas9 knockout, the relatively modest impact of HDAC3is on viral silencing (Figure 5) was surprising. We speculate that HDAC3 may either have an enzymatic role in latency establishment that is incompletely inhibited by chemical inhibition or that HDAC3 may play a non-enzymatic role. Notably, non-enzymatic roles for HDAC3 in transcriptional regulation have been previously reported(H. C. B. Nguyen et al., 2020; Sun et al., 2013). Based on these observations we propose that HDAC1 and HDAC2 together play a crucial catalytic role in viral silencing for which these proteins are individually redundant, while HDAC3 plays an essential but potentially non-catalytic role. Thus, multiple HDACs contribute distinct activities to the process of HIV latency initiation. Further definition of the molecular details of these distinct activities may inform efforts to block viral silencing during reservoir formation and limit reservoir seeding.

## Discussion

The clinical goal of HIV cure research is to achieve long-term remission without therapy in PLWH. The discovery of the slow decay rate of the HIV reservoir(Crooks et al., 2015; Siliciano et al., 2003) has led to efforts to either accelerate reservoir decay through latency reversal or to promote durable proviral silencing(Elsheikh et al., 2019; Margolis et al., 2020; Mediouni et al., 2019). Latency reversal strategies have thus far been hampered by inefficient proviral reactivation(Einkauf et al., 2022; Ho et al., 2013), and this inefficiency limits our ability to clear a broad fraction of the latent reservoir in vivo. Studies show that latent proviruses are likely difficult to reactivate because they are transcriptionally repressed by multiple overlapping mechanisms. Furthermore, latency is likely maintained in vivo via epigenetic mechanisms in the form of histone methylation marks (ie: H3K9me3 or H3K27me3) that allow the repressed viral state to be transmitted during clonal expansion (Spina et al., 2013).

Here, we explore a new approach: targeting chromatin-modifying enzymes that promote initial silencing of transcriptionally active provirus to prevent, rather than reverse, latency. Because, unlike latent viruses, transcriptionally active proviruses are not subject to multiple layers of repression, we propose that preventing latency initiation may be more achievable than latency reversal. For example, blocking the deposition of repressive epigenetic marks at an early stage of infection could prevent key mechanisms of reservoir maintenance. If host cell pathways that regulate entry into latency for a large fraction of infected cells can be identified, then clinical dosing with agents that block these pathways at the time of ART initiation when the majority of the long-lived latent reservoir is stabilized might successfully limit reservoir seeding. By maintaining persistent viral gene expression in productively infected cells in the initial post-ART period, these cells could thus be killed by viral cytopathic effect or rendered vulnerable to immune clearance before transcriptional silencing can occur.

HDACs have previously been shown to repress HIV proviruses during latency, both in in vitro models and in vivo. However, the overall potency of HDACis against latent proviruses in resting CD4 T cells is limited(Spina et al., 2013). We now propose that HDAC activity plays a distinct role in latency *initiation* during effector to memory transition of CD4 T cells, and that histone deacetylation represents a ‘gateway’ event for entry into latency. Our results demonstrate that Class I HDACs play a critical role in initiating latency and that blocking Class I HDAC activity in productively infected CD4 T cells effectively prevents these cells from entering latency during transition to a resting memory state. Furthermore, HDAC inhibition during the transition leads to protracted viral gene expression, even after HDACi removal, suggesting that this treatment reprograms cells into a state that is suboptimal for latency.

By comparing the abundance of various histone marks on the HIV provirus as latency is established, we observed that early HDACi exposure during active infection led to persistently elevated H3K9ac in the gene body and the 3’ LTR. Interestingly, HDACi treatment also led to a substantial reduction in H3K9me3 in the LTR regions. Repressive HMTs like the G9a H3K9 methyltransferase(Imai et al., 2010) require an unmodified lysine residue as a substrate for methylation, and both H3K27me depositing and H3K9me depositing repression complexes can contain Class I HDACs(Imai et al., 2010; van der Vlag and Otte, 1999). Therefore, the locus-specific acetylation of regulatory histone residues can block the addition of these additional repressive and epigenetically transmittable marks, and the reciprocal nature of this regulatory switch likely prevents repression of actively transcribed genes activated by recruitment of histone acetyl transferases like P300. Once repressive methyl marks have been deposited on a provirus and have established robust epigenetic silencing, HDACs likely become less important for proviral repression(Tripathy et al., 2015, p. 27) since HDAC activity in resting cells is low and most proviruses are now in a methylated state. Thus, the fraction of proviruses that lack robust histone methylation, i.e., those that are not “fully silenced” or in “deep latency” will be most responsive to HDACi when used as an LRA. Consistent with this model, longer durations or repeated exposure to HDACi can catch more proviruses in an inducible state(Archin et al., 2017; Shan et al., 2014) likely by tipping the equilibrium towards active proviral gene expression. By contrast, in productively infected activated CD4 T cells, histone acetylation, both globally and at the provirus, is high but is undergoing rapid acetyl group removal by HDACs, leading to a higher sensitivity to HDACis. Genomic experiments like CUT&RUN generate data from an ensemble of cellular genomes and can only identify patterns across a bulk population; thus, novel approaches to link single-cell transcription and chromatin biology may enable a higher resolution picture of proviral regulation and repression(Kaya-Okur et al., 2019; Wu et al., 2021). Functionally, histone repressive marks lead to changes in DNA methylation and nucleosomal remodeling; these mechanisms are important for HIV repression(Arumugam et al., 2021; Conrad et al., 2017; Lu et al., 2022; Rafati et al., 2011) and the respective contribution of each of these factors to latency initiation and maintenance, and their importance to the in vivo latent reservoir, are important questions that should continue to be pursued in HIV cure research.

Our findings also clarify the contribution of individual Class I HDACs toward HIV silencing in CD4 T cells. Specifically, we observed that HDAC3 plays an essential role in latency establishment, but curiously, HDAC3-selective inhibitors had only a minor impact in this system. By contrast, an HDAC1/2i potently blocked latency to a similar degree as vorinostat, together suggesting that the effects of vorinostat are likely mediated through HDAC1 and HDAC2 inhibition but that HDAC3 may also play a separate, non-catalytic role. Thus, we propose that multiple Class I HDACs contribute to HIV silencing in CD4 T cells and that individual HDACs have specialized roles in this process. Identifying the key functional complexes involved in HIV-1 latency initiation may facilitate future advancements in drug specificity, which may provide advantages in developing inhibitors with a robust therapeutic window.

HDAC-regulated acetylation is also a key process in other relevant transcriptional regulatory factors, including the viral protein Tat, the pTEFb transcriptional elongation complex(Bartholomeeusen et al., 2013), and NFKB(Chen et al., 2001; Leus et al., 2016). However, the relative importance of the histone vs non-histone targets of HDACs in HIV latency is unclear. Class I HDACs are abundant nuclear enzymes(Gillespie et al., 2020) but their repressive activity is associated with recruitment to specific genomic loci as members of nuclear co-repressor complexes: the NCOR/SMRT complex for HDAC3 and the NURD, Sin3a, and CoREST complex for HDAC1 and HDAC2(Park and Kim, 2020). Co-repressor complexes are in turn recruited by DNA-binding transcription factors; physical competition can exist between co-activator and co-repressor binding to DNA-binding transcription factors and can be modulated by factors like cell-signaling or ligand binding to steroid hormone receptors (Chen and Evans, 1995; Doetzlhofer et al., 1999; Narita et al., 2003). While a number of mechanisms of HDAC recruitment to the HIV provirus(Margolis, 2011; Romerio et al., 1997; Williams et al., 2006) have been identified in experimental model systems, the extent to which each of these mechanisms operate in primary CD4 T cells remains unclear. Careful functional analysis will be required to assess the relative contribution of HDAC recruitment mechanisms to the HIV provirus, but we anticipate that further advances in our ability to both manipulate and measure the molecular biology of primary human CD4 T cells will empower these future studies. Identification and characterization of repressive mechanisms that are necessary and sufficient for the initiation and maintenance of HIV latency in primary CD4 T cells will inform the future of HIV cure strategies.

Our data also support the conclusion that HIV silencing in CD4 T cells is non-random and occurs preferentially in cells with memory T_SCM_ or T_CM_ phenotypes. The preferential silencing of the provirus in these subsets is related at least in part to the transcription factor FOXO1(Roux et al., 2019, p. 1; Vallejo-Gracia et al., 2020, p. 1); preferential silencing in memory T_SCM_ and T_CM_ cells coupled to the longevity and proliferative capacity of these cells is a major factor in the persistence of the latent reservoir. Cells with these phenotypes are preferentially found in clonal expanded cell populations with HIV proviruses(R. Liu et al., 2020; Maldarelli et al., 2014; Murray et al., 2016; Simonetti et al., 2021). Interestingly, we observe that HDACi exposure during effector to memory transition for HIV infected cells leads to a reprogramming of the subset identity, including a dramatic reduction in abundance of the highly stable and self-renewing T_SCM_ cell subtype. These cells are hypothesized to represent a key component of the long-lived clinical reservoir(Buzon et al., 2014). Thus, HDACis may be effective in vivo at preventing seeding of HIV latency within this subset, potentially leading to a less durable or self-renewing reservoir. The association between the latency prevention activity of HDACis and their impact on CD4 T cell subset differentiation will need to be investigated further.

In this study, we demonstrate that HDACis potently maintain viral gene expression from proviruses that are already transcriptionally active. The ability of HDACis to prevent HIV latency when administered to actively infected cells suggests a novel clinical approach to limiting the size of the HIV reservoir. Since a large fraction (∼70%) of the reservoir originates from infection events near the time of ART initiation(Abrahams et al., 2019). As such, it may be possible to co-administer HDACis at the time of ART initiation to prevent actively infected cells from transitioning to the long-lived latent state. Extending viral gene expression in these cells may, in turn, be sufficient to lead to elimination of these cells through cytopathic effect or by immune clearance. Although the relative contribution of these trajectories to seeding the latent reservoir in vivo is unclear, it is notable that ART initiation is accompanied by a large redistribution of CD4 T cells from effector to memory subsets(Lempicki et al., 2000; Mohri et al., 2001). An important consideration is the level of viral gene expression required for clearance by specific viral/host mechanisms. It remains unclear how a certain level of viral gene expression affects the likelihood of clearance, but this will be an important parameter that will need to be careful defined in the HIV latency and persistence field. However, our studies raise testable hypotheses about the role of HDACs in the formation of the HIV reservoir; thus, testing these hypotheses with in vivo models and clinical studies should provide important mechanistic and therapeutic insights into reservoir formation.

We acknowledge several important limitations when considering this work. First, the in vitro system here does not necessarily reflect the all the mechanisms responsible for the generation of all latently infected cells in vivo. While we model activated T cells returning to a resting state, it is likely that some proviruses are silenced immediately after integration and do not experience a period of productive viral gene expression. Thus the success of targeting latency initiation may depend on the fraction of the reservoir for which a given intervention can prevent further latency enforcement versus the fraction that are already sufficiently repressed to allow persistence(Spina et al., 1997, 1995; Swiggard et al., 2005). Furthermore, the activation conditions of our T cell culture, CD3/TCR cross-linking under high IL-2/IL-7 conditions, may be a stronger activation stimulus than encountered by most cell in vivo, and the strength of TCR stimulation has previously been linked to the likelihood of the HIV provirus entering latency(Gagne et al., 2019). Finally, we note the use of a non-replication competent virus that is missing several accessory proteins which could affect important aspects of latency initiation such as cell-survival and potentially transcriptional regulation.

In summary, these findings highlight the key role of HDACs as initiators of viral entry into latency and indicate that HDACis could be used to trap HIV in infected cells in a state of active viral gene expression. Furthermore, we observe that this activity acts on both the provirus to promote an acetyl/methyl switch on H3K9 residues within proviral histones and on the host-cell to promote a non-proliferative cell-state. Further basic science exploration of latency initiation and the clinical application of these insights may enable new approaches towards an HIV cure.

## Materials and Methods

### STAR METHODS

#### RESOURCE AVAILABILITY

##### Lead Contact

Further information and requests for resources and reagents should be directed to and will be fulfilled by the lead contact, Edward Browne.

##### Material availability

HIV-dreGFP/Thy1.2 will be deposited to Addgene plasmid repository. Other materials employed in this study are available commercially.

##### Data and code availability

Next-generation sequencing data from CUT&RUN and RNA-seq experiments have been deposited to NCBI GEO and will be available pending completion of publication process. Other data are available upon reasonable request to the corresponding author.

## EXPERIMENTAL MODELS AND SUBJECT DETAILS

Healthy (HIV-negative, Hepatitis B-negative) donor blood products were acquired from leukapheresis reduction chambers from apheresis platelet donations. The collection of these products was performed under an Institutional Review Board by Stemcell Technologies and purchased by our laboratory commercially. Review by the University of North Carolina IRB determined that the studies described herein constitute analysis of secondary samples derived from human subjects and were thus deemed exempt from IRB review.

## METHOD DETAILS

### Primary CD4 T-cell infection model

CD4 T cells were obtained by gradient centrifugation and magnetic isolation from leuko-reduction blood products (StemCell Technologies). Blood products were washed and centrifuged on a Ficoll gradient to obtain peripheral blood mononuclear cells (PBMCs). PBMCs were treated with ACK lysis buffer to remove residual red blood cells, and CD4 T cells were isolated from PBMCs with a negative magnetic isolation kit (StemCell Technologies) and examined for purity using anti-CD3 and anti-CD4 flow-cytometric staining. Cells were then either immediately cultured or frozen in cryopreservation media composed of 90% Fetal Bovine Serum and 10% DMSO. After isolation or thawing, cells were activated using anti-CD3/anti-CD28 Dynabeads (Gibco) in RPMI R10 (10% FCS; 1% Penicillin Streptomycin; 10mM HEPES; 2mM L-Glutamine; 1mM Sodium Pyruvate) supplemented with 100U/mL IL-2 and 1ng/mL IL-7 (Peprotech). After 48h of stimulation, cells were thoroughly resuspended and activating beads were removed. Activated CD4 T cells were resuspended in viral supernatant (see below) containing 4μg/mL polybrene (Hexadimethrine Bromide), and cells were spinoculated in 50mL conical flasks in 5mL aliquots at a density of 10^6^ cells/mL at room temperature for 2h at 600RCF. Following spinoculation, viral supernatant was removed and replaced with RPMI R10 at 10^6^ cells/mL. 48-72h after spinoculation, cells were examined by flow cytometry to identify the percent infected cells as measured by GFP and Thy1.2 expression. In some experiments infected cells were isolated by Thy1.2 surface marker magnetic-bead based enrichment (Stemcell Technologies) and in other experiments were cultured without magnetic isolation using Thy1.2 staining as a marker for infected cells. Media was replaced every 2-3 days with complete RPMI supplemented with recombinant 100U/mL IL-2 and 1ng/mL IL-7 at 10^6^ cells/mL. After 7d of cell expansion, cell density was increased to 2×10^6^ cells/mL. Viral gene expression was measured weekly by flow cytometry for GFP and Thy1.2, with the fraction of infected (Thy1.2+) cells expressing detectable GFP measured using uninfected control cells to set a GFP+ gate. Latency re-initiation was examined by infecting cells as above and maintaining cells in culture for 21d. After 21d, cultures were examined by flow cytometry to confirm low viral gene expression and subsequently restimulated for 48 hours using anti-CD3/anti-CD28 coated Dynabeads. Reactivated viral gene expression was confirmed by flow cytometry for Thy1.2 and GFP at 48h, and cellular activation was confirmed by tracking cellular proliferation.

### Flow cytometry

Flow cytometry for productive viral gene expression was performed as described in text; infected cells were prepared for flow cytometry following PBS washes, Live-Dead staining (Zombie Violet, Biolegend 1:1000), and Thy1.2 staining (1:1000). Samples were fixed in 4% PFA for 10 minutes and analyzed immediately or stored overnight at 4° C. For CD4+ memory marker flow cytometry, cells were stained with Live-Dead viability dye (Zombie NIR), followed by antibodies to CD3, CD4, CD8, CD45RA, CCR7, CD28, CD95, CD127a, PD-1, CD90.2, CD14, CD16, CD19, and CD56.

Samples were acquired on a Fortessa flow cytometer (Becton Dickson) and analyzed using FlowJo (version ×10.0.7r2). At least 10,000 CD4 T cells events were collected for analysis. Compensation controls were prepared for each fluorochrome using anti-mouse Ig, κ compensation particles or anti-rat Ig, κ(CD90.2). For phenotypic markers (CD45RA, CCR7, CD28, CD95, CD127a, PD-1, CD90.2), positive events were gated using fluorescence minus one controls.

### Small-molecule inhibitor studies

Inhibitors (see key resource table below) were added to culture media for the indicated time periods and refreshed three times weekly, with the exception of experiments described in figure 1H, which involved twice daily media changes. Drug dilutions were prepared by serial dilution in DMSO or using the D300e Digital Dispenser (Hewlett-Packard). Final DMSO concentrations in culture media were maintained at or below 0.1%. Flow cytometry data were collected and analyzed as above in Flowjo and fit to a 4-parameter Hill equation in Graphpad Prism.

### Cellular fractionation and western blotting

To produce whole cell lysates, cells were harvested, washed in 1x phosphate buffered saline (PBS), and lysed in RIPA buffer (ThermoFisher) supplemented with the 1x protease inhibitors (Roche) 100μM of the pan-HDACi Trichostatin A (TSA) and 1% Pierce Universal Nuclease (ThermoFisher). Cell fractionation was performed with a commercially available kit (ThermoFisher). Sequential lysis was performed to isolate fractions containing 1) soluble cytoplasmic proteins through hypotonic lysis, 2) membrane-bound proteins, 3) soluble nuclear proteins, and 4) DNA-bound proteins liberated by treatment with micrococcal nuclease. To compensate for the small size of CD4 T cells compared with commonly used cell-lines, 2×10^6^ cells per condition were lysed in volumes recommended for 1×10^6^ cells. Fractionated cell lysates were quantified by Bradford assay (ThermoFisher), resolved by Tris-Glycine SDS-PAGE, and transferred to polyvinylidene difluoride (PVDF) or nitrocellulose membranes for the detection of transcription factors or histones respectively. Blots were blocked in 5% milk in 1x Tris Buffered Saline (TBS), probed overnight with antibodies diluted in TBS containing 5% BSA or 5% milk, and washed with 0.1% Tween20 TBS (TBST). Membranes were probed with secondary antibodies (1:10,000) conjugated to horseradish peroxidase for 1h at room temperature and washed in TBST, followed by treatment and detection of luminescence by enhanced chemiluminescence (ThermoFisher).

### CUT&RUN assays

CUT&RUN was performed with a commercially available kit according to manufacturer instructions with some modifications (Epicypher). Healthy donor T cells were infected with HIV-dreGFP/Thy1.2 and isolated based on Thy1.2 surface expression. Infected cells were cultured for 21d, during which cells exhibited progressive decline in viral gene expression as latency is established. Latently infected cells were then re-stimulated with anti-CD3/anti-CD28 dynabeads to increase cell numbers and then allowed to return to latency. As the reactivated cells returned to a resting state, cells were treated in triplicate with or without vorinostat (335nM) for 21d. 3×10^5^ CD4 T cells per antibody condition were harvested, washed, and coupled to Concanavalin A-coated beads for 30 min at 4° C with mixing. Cells were incubated overnight in Digitonin-containing (0.01%) antibody buffer with the indicated antibodies at the following concentrations: IgG 1:50, H3K4me3 1:50, H3K27ac 1:100, H3K27me3 1:17, H3K9ac 1:17, H3K9me3 1:80, RNA pol2 1:17, RNApol2 P-S2 1:50. After incubation, cells were washed twice in cell permeabilization buffer (buffer CP, 0.01% Digitonin) and incubated with pAG-MNase for 15 min at room temperature. Cells were washed twice in buffer CP before CaCl_2_ was added for a 2h cutting reaction at 4°C with rotation. Stop buffer containing EDTA, RNase A, and *E. coli* spike-in DNA was added to halt the reaction during a 37° 10-min incubation step. Supernatants were collected and added to DNA binding buffer for subsequent DNA isolation by spin columns. Supernatant DNA yields were eluted from columns in 12μL of Tris-HCl elution buffer and quantified by Qubit High-Sensitivity dsDNA assay. CUT&RUN Next-Generation Sequencing (NGS) library preparation was performed using NEBNext Ultra II DNA library preparation kits (New England Biolabs). DNA end repair and A-tailing was performed followed by adapter ligation to NEBNext hairpin adaptors and linearization of NEBNext adaptors with USER enzyme. Adaptor ligated libraries were amplified with sample-specific barcoded dual-index primers with PCR conditions optimized for amplification of short fragments: 14 cycles 15s at 98°C and 10s at 60^°^C. Libraries were quantified and examined by Qubit dsDNA assay (ThermoFisher) and capillary electrophoresis (Agilent Tapestation). Quantified libraries were pooled and subjected to paired-end sequencing with the NovaSeq 6000 platform (Illumina) at an approximate read-depth of 20M read-pairs/sample. Analysis was performed as follows: FASTQ files were demultiplexed according to sample-index using the mkfastq command. Reads were filtered for adapter contamination using Cutadapt(Martin, 2011) and filtered such that at least 90% of bases of each read had a quality score>20. Duplicated sequences were then capped at a maximum of five occurrences, and reads were aligned to a custom reference genome that concatenated GRCh38 (hg38) with the HIV-dreGFP/Thy1.2 sequence and *E. coli* K12 sequence each added as additional pseudo-chromosomes. Alignment was performed using STAR(Dobin et al., 2013) version 2.7.3a retaining only primary alignments. Reads overlapping regions of the genome deemed problematic by ENCODE were excluded. Regions of significant enrichment were identified using MACS2(Zhang et al., 2008) with a threshold of q < 0.1, and H3K27me3 data were run in “broad” mode. A union set of peaks called for all samples and conditions was then assembled using BEDtools(Quinlan and Hall, 2010). Regions exhibiting significant changes in abundance between vorinostat and vehicle-control conditions were detected using CSAW(Lun and Smyth, 2016) wherein the tested windows (10kb for H3K27me3, 250 bp for all others) were restricted to those overlapping the union peak set, and significance was reported if combined p < 0.01. Signal over promoters was computed with deepTools(Ramírez et al., 2016, p. 2) multiBigWigSummary with GENCODE v36 transcription start sites +/- 500 bp.

### mRNA-sequencing

Media containing 500nM Vorinostat or 1:1000 DMSO was replenished three times weekly for 21 days. RNA was extracted from 10^5 CD4 T cells using the RNeasy kit (Qiagen, #74134). cDNA synthesis was accomplished using the SMART-HTv4 (Takara, #634888) involving cDNA synthesis and cDNA amplification (7 cycles) using a template-switching oligo. Illumina sequencing libraries were prepared using the Illumina Nextera XT kit (Illumina #FC-131-1024) to generate tagmented cDNA followed by indexing PCR with a dual-index strategy. Sequencing libraries were quantified and examined by Qubit dsDNA assay (ThermoFisher) and capillary electrophoresis (Agilent Tapestation) before pooling at an equimolar ratio and sequencing by HISEQ4000 HO 2×50 PE sequencing (Illumina). Sequencing quality reports were generated with fastQC v 0.11. Sequencing adapters were trimmed from 3’ ends of reads using cutadapt v2.9 and FASTX toolkit v0.0.14 was used to identify high-quality reads with options -q 20 -p 90. Reads were aligned to the GRCh38 human genome using STAR v2.7.7a with options: --quantMode TranscriptomeSAM GeneCounts --twopassMode Basic --outFilterMismatchNmax 2 --alignIntronMax 1000000 -- alignIntronMin 20 --chimSegmentMin 15 --chimJunctionOverhangMin 15 --outSAMtype BAM Unsorted --outFilterType BySJout --outFilterScoreMin 1 --outFilterMultimapNmax 1. Expression of genes was quantified from the GENCODE v38 comprehensive gene annotation set restricted to reference chromosomes (gencode.v38.annotation.gtf.gz) using htseq-count v0.6.17 with options -f bam -r name -s no -a 10 -t exon -i gene_id -m intersection-nonempty. Differential expression testing was accomplished in R using DESeq2 v1.34.0. For alignment of HIV reads, a custom HIV reference genome containing the HIV-dreGFP/Thy1.2 sequence was constructed using STAR v2.7.7a using the genomeGenerate command. RNAseq reads were then aligned to the HIV genome using STAR v2.7.7a with the option --genomeSAindexNbases 5 to account for the small reference genome size.

### CRISPR-CAS9 gene knockout studies

Protospacer targeting sequences against HDACs were pre-designed by Integrated DNA Technologies (Key resource table), while Tat targeting and non-targeting control sequences were derived from previous literature(Ophinni et al., 2018; Ting et al., 2018). CRISPR-CAS9 ribonucleoprotein complexes (RNPs) were prepared as follows: crRNA targeting sequence and tracrRNA (IDT) were mixed 1:1 at a final concentration of 100μM and annealed in duplex buffer (IDT) by heating to 95°C and slow cooling to room temperature in a PCR thermocycler. RNP complexes were prepared the day of nucleofection by mixing 0.8μL of CAS9 enzyme (62μM concentration, 49.6pMol total, ALT-R by IDT), 0.8μL of poly-glutamic acid(D. N. Nguyen et al., 2020) (15KDa average polymer size, 100mg/mL concentration, Alamanda laboratories), and 1μL of annealed duplex crRNA:tracrRNA (100pMol for a final molar ratio of approximately 2:1 RNA:CAS9 enzyme). Effective targeting sequences were identified in pilot experiments, and targeting of HDAC3 was multiplexed for more efficient target knockout as described previously(Oh et al., 2019; Seki and Rutz, 2018) and as specified in key resource table. Primary CD4 T cells were isolated, activated, and infected as above. Six days after activation (4dpi), 3×10^6^ cells per condition were pelleted at 90 RCF for 10 minutes, washed once in PBS, and resuspended in primary cell nucleofection buffer P3 (Lonza) at a concentration of 1×10^8^ cells per mL (2×10^6^ cells per 20μL reaction volume). Cell suspensions were added to RNP complexes, and nucleofection of T cells with RNP complexes was accomplished using the CM137 protocol (Lonza 4D nucleofector device) based on a published protocol in human T cells(Ting et al., 2018). Following nucleofection, cells were immediately resuspended in pre-warmed complete RPMI and incubated at 37° for 20 min prior to resuspension at a density of 2×10^6^ cells per mL in complete RPMI supplemented with IL-2 and IL-7 as above. Viral gene expression was examined as above at the indicated time points. Seven days after RNP nucleofection, 2×10^6^ cells were collected for western blot analysis.

### Plasmids and viruses

HIV-dreGFP/Thy1.2 was generated by cloning an IRES-Thy1.2 expression cassette into an NL4-3-Δ6-dreGFP reporter virus(Yang et al., 2009), immediately 3’ to the stop codon of the dreGFP gene in the NheI restriction site. Lentiviral production was accomplished using a 3-vector transfection system, with Gag and Pol supplied by psPAX2 and envelope pseudotyping with the vesicular stomatitis virus glycoprotein (VSV-g). 293T/17 cells (ATCC) were transfected with HIV-dreGFP/Thy1.2, pMD2-VSVg, and psPAX2 using the Mirus LT1 transfection reagent at a 3:1 volume to mass ratio. After 24h, DMEM was replaced with RPMI R10. After 48h, virus-containing supernatant was cleared by centrifugation at low speed and filtered through a 0.45μm low protein-binding filter. Viral supernatants were used immediately for spinoculation of primary T cells.

## QUANTIFICATION AND STATISTICAL ANALYSIS

Statistical tests and significance cutoffs are described in figure legends and expanded upon below. Statistical analyses were performed in Graphpad Prism or R. Statistical analyses for CUT&RUN and RNA-seq experiments followed field standard methods including negative binomial peak calling with MACS and DEG testing using EDGER; further details are discussed in the respective methods sections. No assumptions of normal data distributions were made for statistical difference testing of flow-cytometry data and instead a Wilcoxon ranked-sum test was employed; in the case of flow-cytometry experiments statistical testing was not attempted when studies were assessed as underpowered for a Wilcoxon ranked-sum test. Statistical analysis was facilitated by the UNC Center for AIDS Research biostatistics core.

The mixed effects model employed in Figure 1B follows the form *Yij=β0+tijβ1+bi+єij*, where *t*ij*,Y*ij denote the time (in days) and the %GFP(+) for subject *i* at time *j*, respectively; and β0 be and β1 are the two fixed effects referring the mean %GFP(+) at zero days and the change rate of %GFP(+) over time, respectively. The random effects include *bi* for modeling the heterogeneity of %GFP(+) for donor *i*, and є, a random error term. We assume a compound symmetry covariance structure, that the random effects *bi* are independent and identically distributed (i.i.d.) with distribution *N*(0,*σb*2), and that the errors are i.i.d. with distribution *N*(0,*σb*2). This mixed effect model assumes that different donors have the same slope (i.e., the same rate of change of %GFP(+) with respect to time). The estimated fixed effect of time on value is β1=-2.5789 (t = −10.67 on 23 degrees of freedom, p < 0.0001), or on average, %GFP+ of Thy1.2+ decreases by −2.5789% per day.

Figure 4C employs a generalized estimating equation (GEE) to test if HIV-aligning reads per 10^6^ reads differs between the DMSO and vorinostat treated samples. We assume the data follow a Poisson distribution and use a log link function in the GEE model. The parameter estimate for the effect of Vor on the outcome is *βvor*=0.7713, with a Wald χ^2^=15153.8, and p-value <0.0001. We interpret this to mean that the log of the mean count of HIV-aligning reads in the Vor group greater than the log of the mean in the DMSO group by 0.7713. Equivalently, the mean count of HIV-aligning reads in the Vorinostat treatment group is *e*0.7713=2.163 times greater than the mean count in the DMSO group.

## KEY RESOURCES TABLE

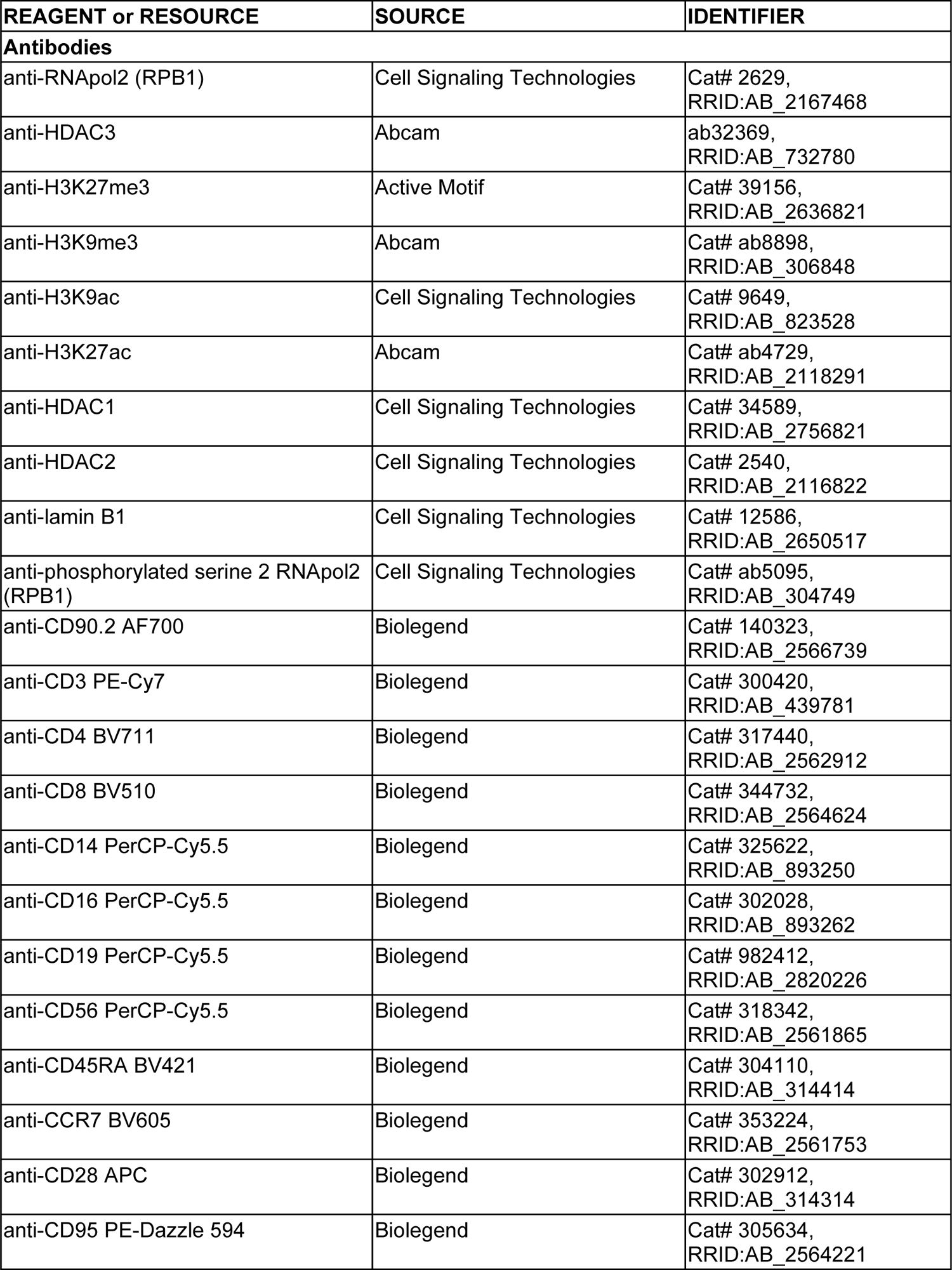
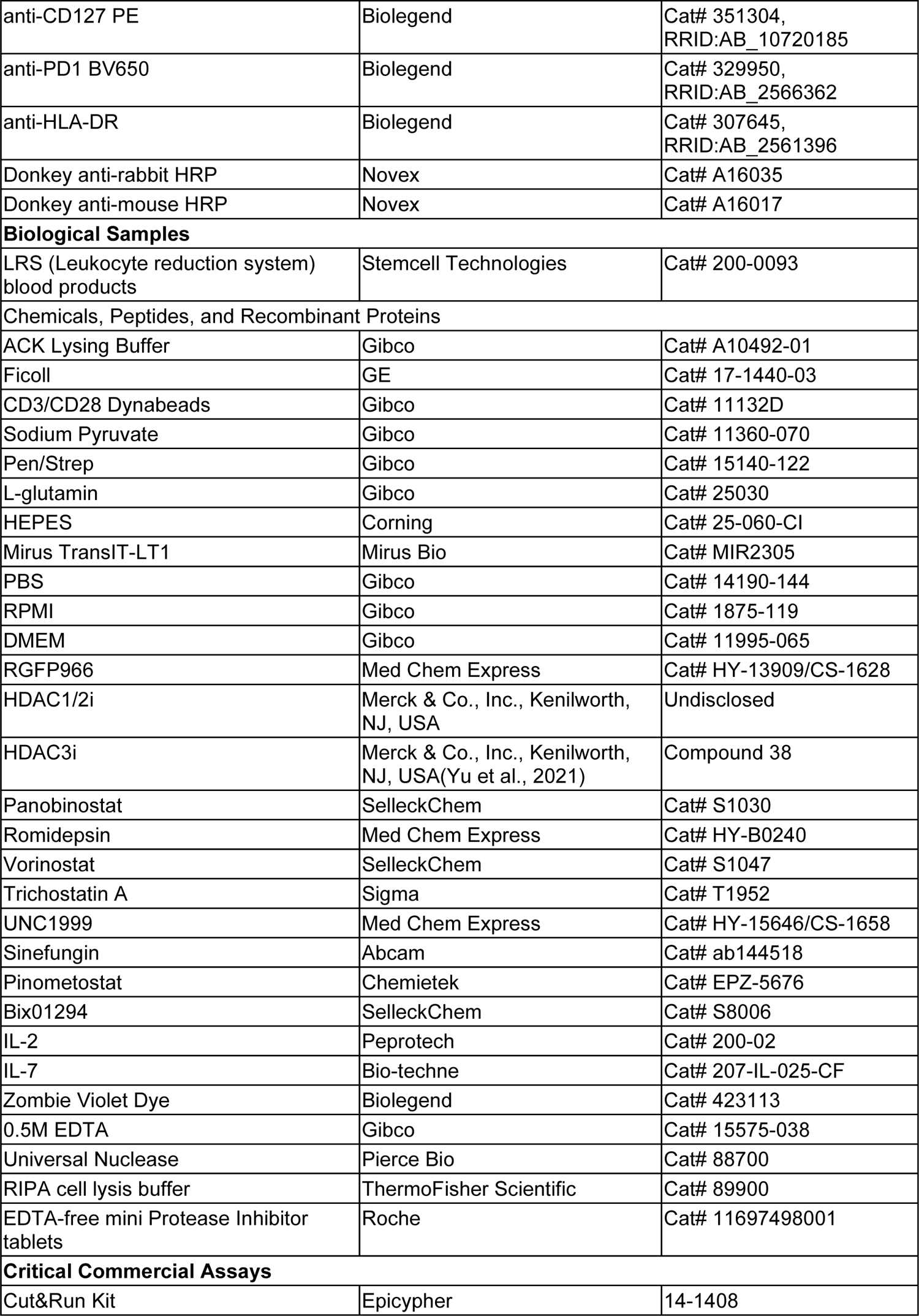
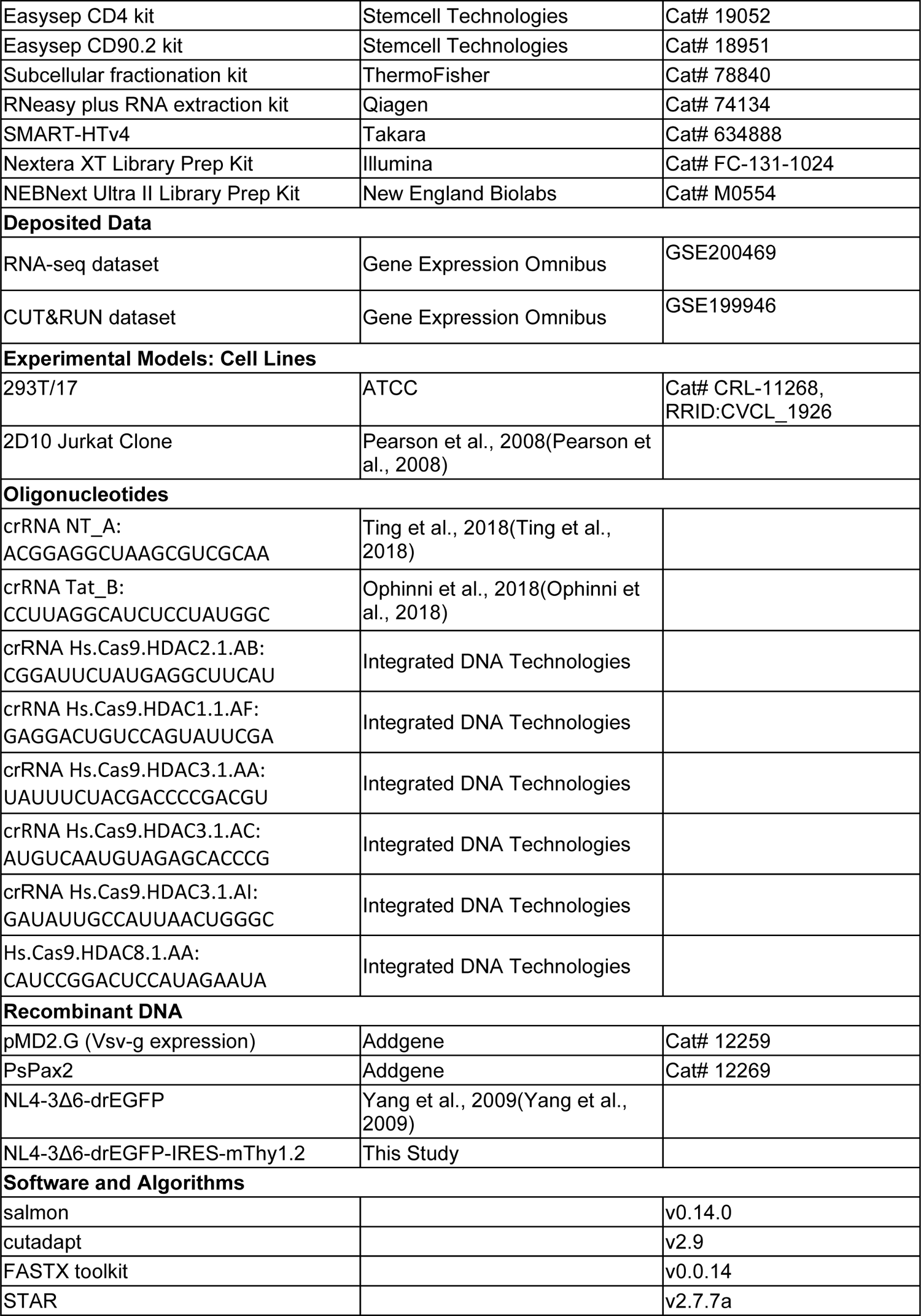
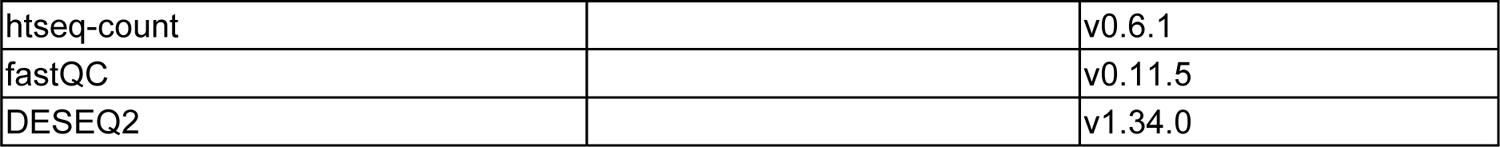

## Acknowledgements

HDAC1/2i and HDAC3i were provided by Merck & Co., Inc., Kenilworth, NJ, USA. Next-generation sequencing studies were facilitated by the UNC School of Medicine High-Throughput Sequencing Facility. Analysis of CUT&RUN data was facilitated by the UNC School of Medicine Bioinformatics and Analytics Research Collaborative (BARC). Statistical analysis was facilitated by the UNC Center for AIDS Research Biostatistics Core. This work was supported by the following grants from the National institutes of Health: NIAID #5-R01AI143381-03, NIAID #5-UM1AI164567, NIDA # 5-R61DA047023-03, NIAID #5-T32AI007419-28 (UNC-Chapel Hill Molecular Biology of Viral Diseases T32), and funding from Qura therapeutics.

## Contributions

J.J.P. and E.P.B. conceptualized the study. D.M.M., N.G., and E.P.B. oversaw the research. J.J.P., C.A.L., S.D.B., A.M., Y.X., and A.A.R. performed experiments. G.C., B.R., F.Z., and J.M.S. assisted in data analysis. J.J.P. and E.P.B. wrote the first draft manuscript and all authors contributed to subsequent drafts.

## Supplemental information

**Supplemental figure S1.**
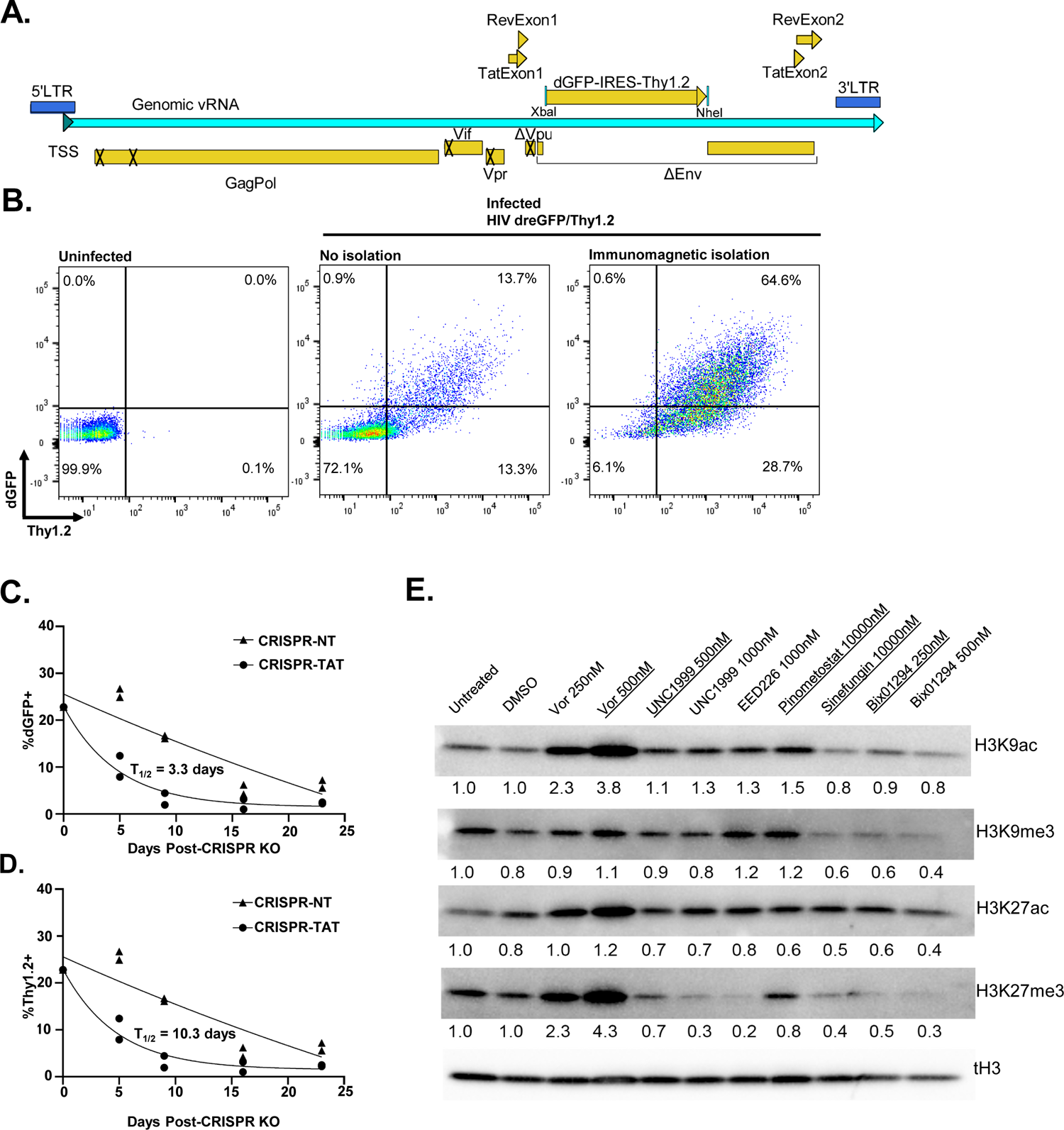
**A.** Schematic of HIV-dreGFP/Thy1.2 reporter virus. Diagram is to approximate scale, 10.5kb total length. Sequence available in Supplementary File 1. **B.** Immunomagnetic isolation of CD4 T cells infected with HIV-dreGFP/Thy1.2. Uninfected (left) or infected cells (right two panels) were analyzed by flow cytometry for GFP and Thy1.2 expression before and after immunomagnetic enrichment of Thy1.2+ cells. **C,D.** Resting infected CD4 T cells were activated by anti-CD3/CD28 beads and nucleofected with sgRNA/Cas9 RNP complexes directed against viral Tat or with a non-targeting control. The frequency of GFP+ and Thy1.2+ cells was measured by flow cytometry, and data were fit to a one-phase decay model in Graphpad Prism. **E.** Immunoblot of histone epigenetic marks in 2D10 Jurkat cells treated for 72h with the indicated inhibitors. Membranes were stripped with a low-pH stripping buffer, and signal from histone post-translational marks was normalized to total histone H3 signal re-probed on the same membrane. Underlined conditions indicate doses employed in experiment in Figure 1D.

**Supplemental figure S2.**
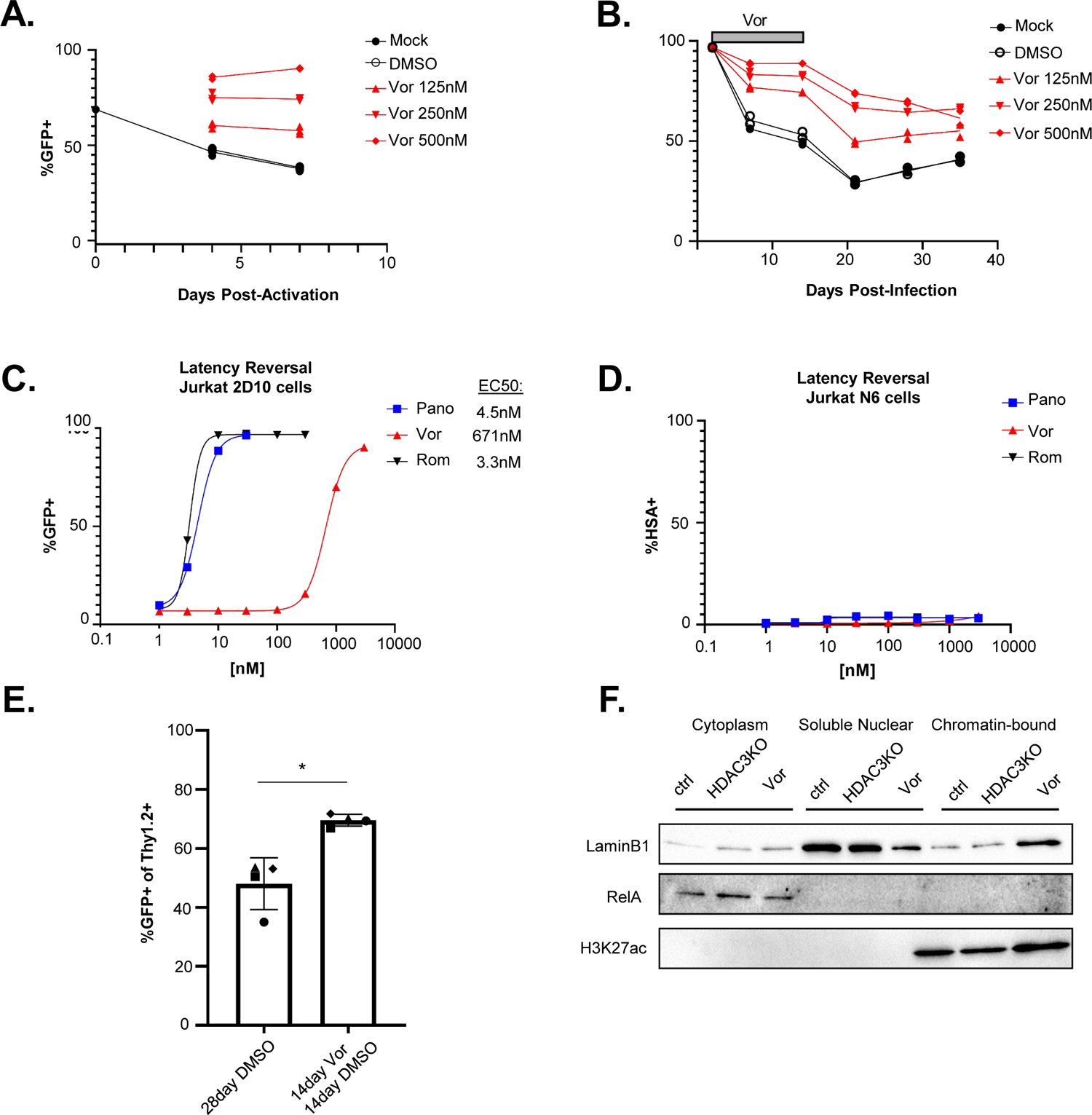
**A.** Latently infected cells were isolated by flow-sorting and cultured for 8 weeks using the H80-feeder cell culture model(Bradley et al., 2018; Sahu et al., 2006), re-stimulated by anti-CD3/CD28 beads to reverse latency, then treated with the indicated doses of vorinostat. Re-entry into latency was measured by flow cytometry for GFP expression. **B.** HIV-dreGFP/Thy1.2-infected cells at 2dpi were treated with the indicated doses of vorinostat or vehicle control (DMSO), and the frequency of GFP+ cells was measured by flow cytometry for 14d. After two weeks in the presence of DMSO vehicle or vorinostat, vorinostat was removed from the media and the cells were then cultured for an additional 14d. **C,D**. Dose response curves of latency reversal with 48h HDACi exposure in 2D10 (C) and N6 (D) Jurkat clones. Viral reporter expression was measured with flow cytometry and fit with a non-linear Hill model. **E.** The percentage of GFP+ cells in the infected cell (Thy1.2) population at 28dpi was quantified after 14d with vorinostat followed by 14d without or after 28d with control vehicle (DMSO) only. Data shown are derived from two independent experiments using CD4 T cells from four independent donors. Statistical comparison using the Wilcoxon ranked sum test; *=p<0.05. **F.** Immunoblot in resting CD4 T cells after fractionation into cytoplasmic, soluble nuclear, and chromatin-bound protein fractions.

**Supplemental figure S3.**
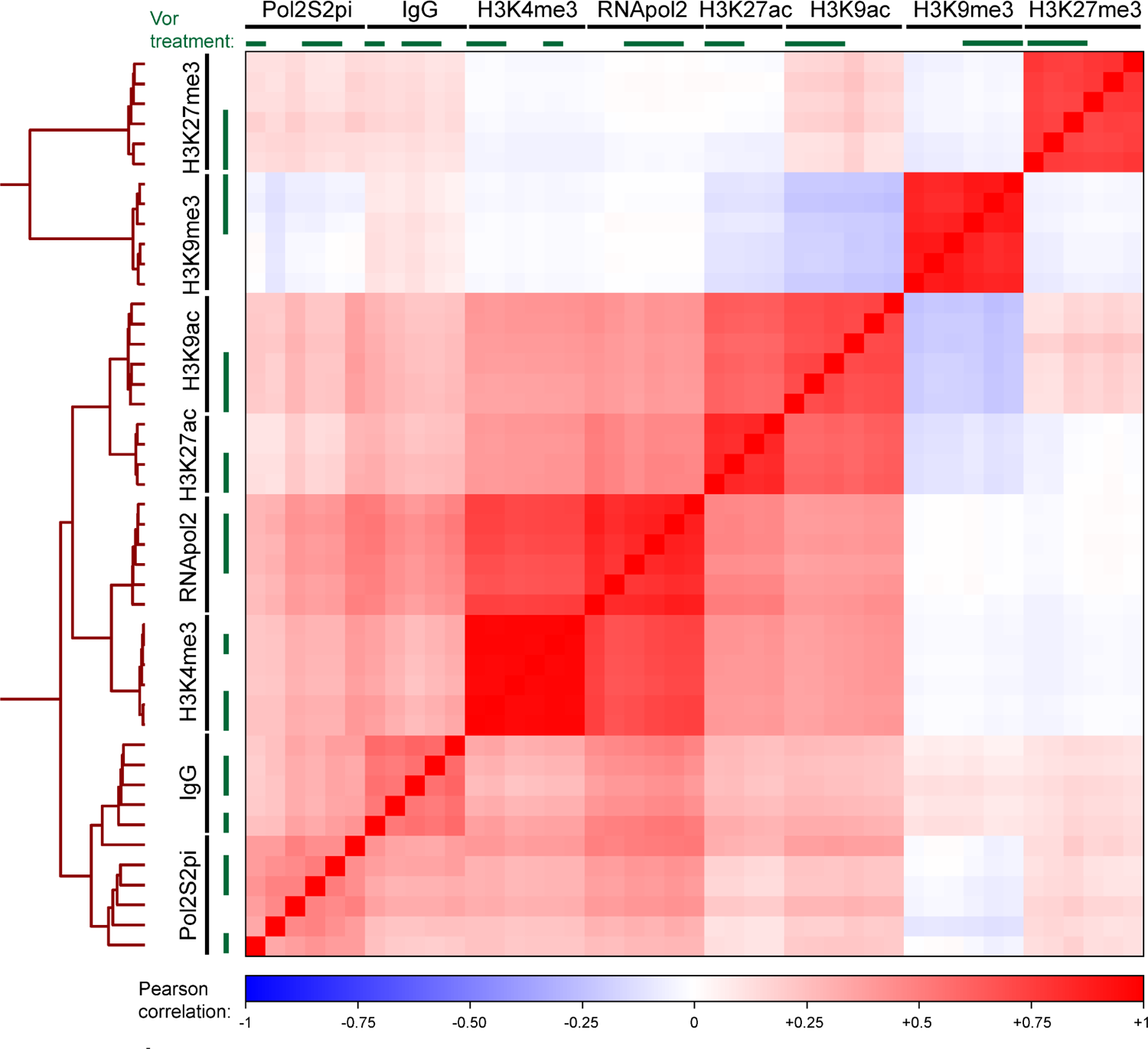
Pearson correlation matrix of CUT&RUN data analysis. Pearson correlation coefficients from each sample over the union set of peaks was called across all factors and conditions. The signals were computed and plotted using deeptools multiBigWigSummary and plotCorrelation. Correlation values were plotted on a scale of −1 (blue) to 0 (white) to +1 (red)

**Supplemental figure S4.**
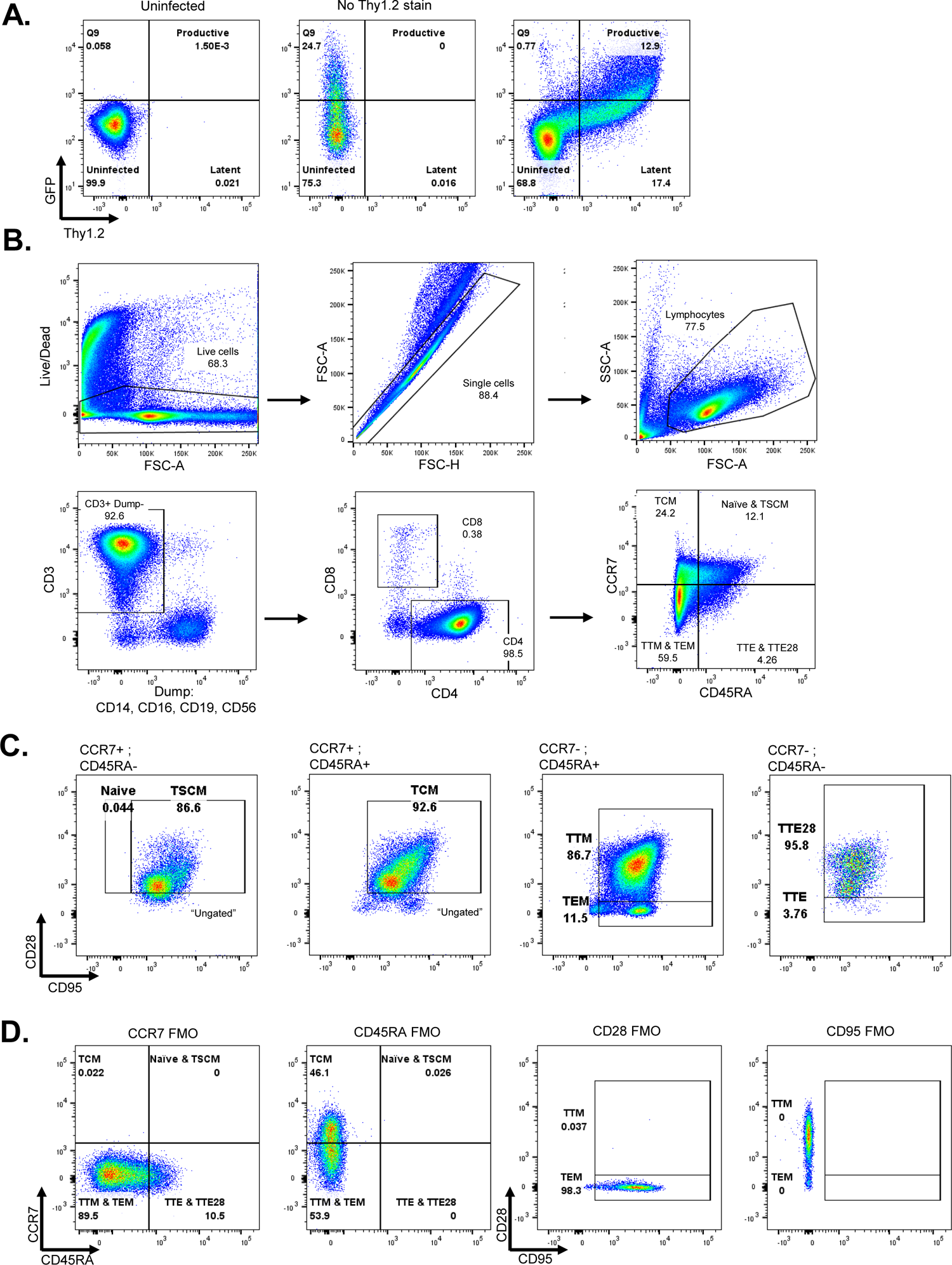
**A-D.** Gating strategy for primary human CD4 T cells in multiparameter flow cytometry analysis. Compensation was performed in Flowjo using standard matrix-adjusted practices based on single-color stained controls. Gating was performed in Flowjo based on Fluorescence-minus-one controls depicted in (**D**).

**Supplemental figure S5:**
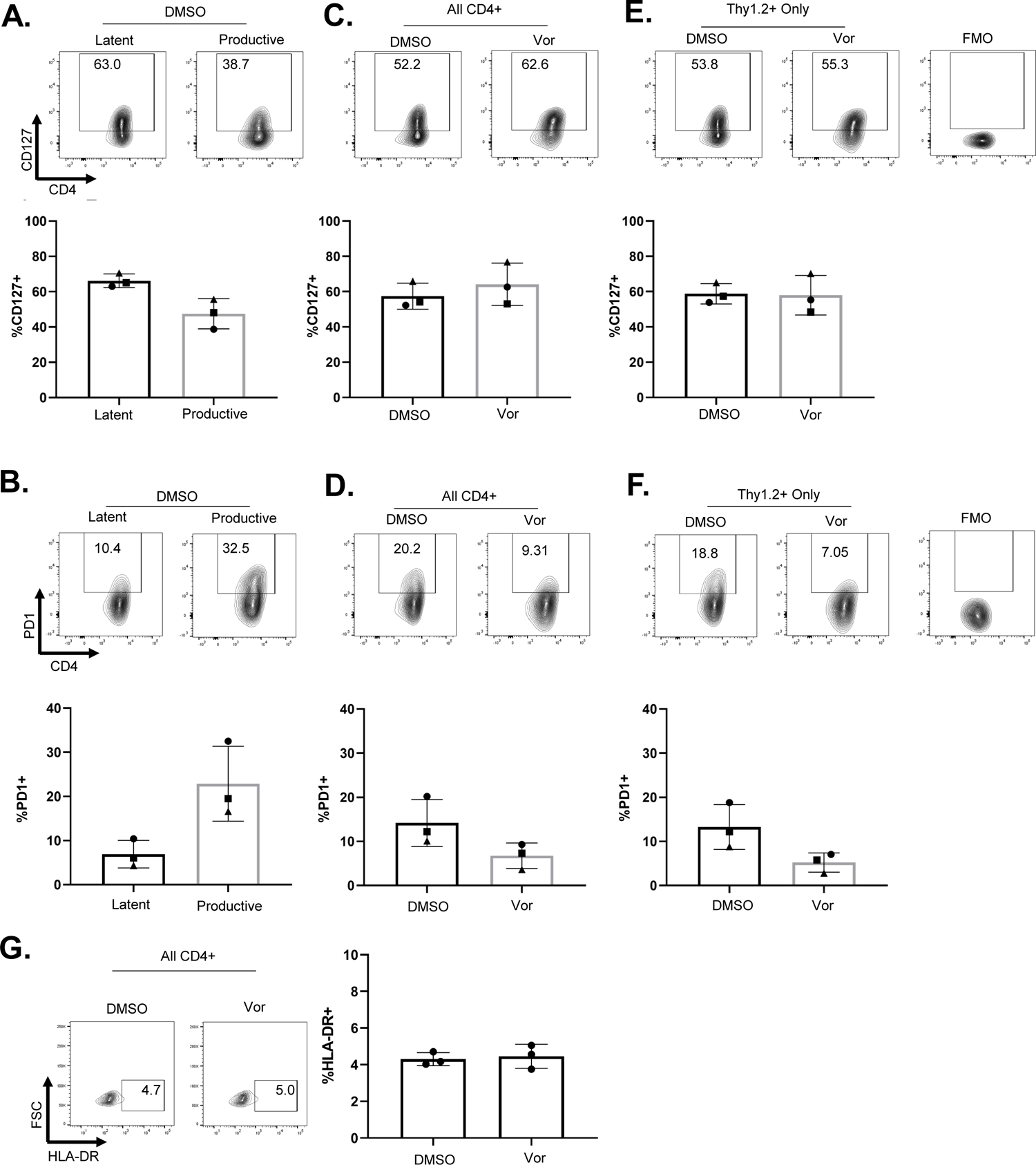
**A-G.** Top: Representative flow plots of the indicated surface markers. **A-G** Bottom: Frequency of cells expressing the indicated markers in DMSO treated cells split into latent and productive subsets (**A,B**) or split between vehicle- and vorinostat-(**C,D,E,F**) treated cells. Bars represent mean and standard deviation of three CD4 T cell donors analyzed in parallel 14dpi (**A-F**) or one cell-donor analyzed in triplicate 21dpi (**G**). Error bars represent standard deviation. **C,D.** Gating strategy for CD127+ and PD-1+ CD4+ cells. **E,F**. Gating performed only on infected (Thy1.2+) cells in culture.

**Supplemental figure S6:**
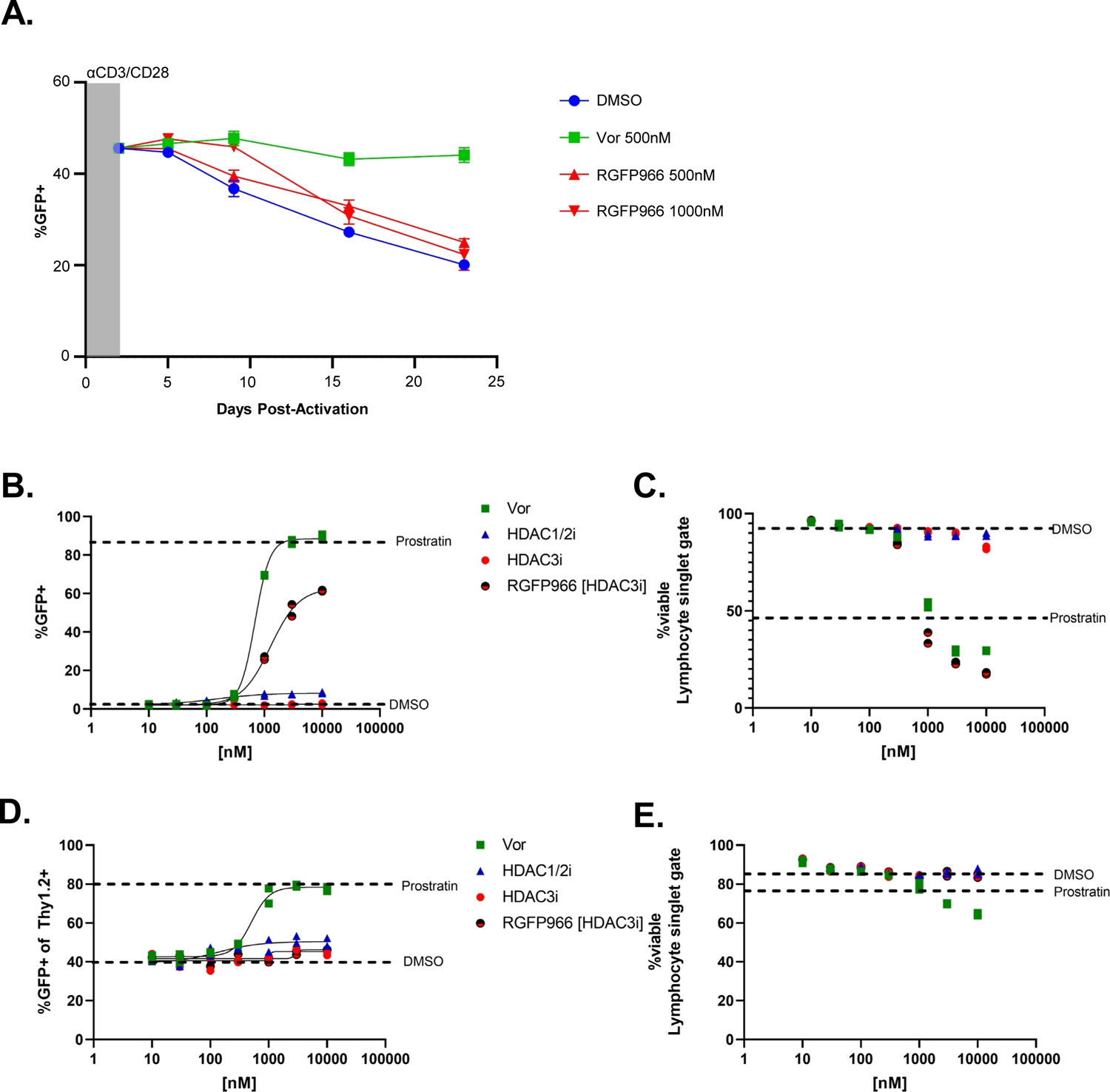
**A**. Latency re-initiation experiment with a structurally distinct HDAC3i. Latently infected primary T cells selected by flow cytometry sorting were stimulated with anti-CD3/CD28 beads and cultured in the presence of Vor (500nM) or the HDAC3 inhibitor RGFP966(500nM or 1000nM). **B-E**. Latency reversal experiments after 48-hour treatment with a dose-response curve of the indicated inhibitors. Two technical replicates were collected and fit to a four parameter Hill model. **B**. Latency reversal in 2D10 cells was measured by flow cytometry of cells across a three-fold dilution series of inhibitor doses. **C**. Viability of cells at each inhibitor dose as measured by Zombie Violet membrane exclusion dye in the lymphocyte->singlet gate. **D**. Latency reversal in infected primary T cells was determined by measuring the frequency of Thy1.2+;GFP+ cells across an inhibitor dose response curve as in (**B**). **E**. Viability of cells plotted as in (**C**).

## References

1. Abrahams, M.-R., Joseph, S.B., Garrett, N., Tyers, L., Moeser, M., Archin, N., Council, O.D., Matten, D., Zhou, S., Doolabh, D., Anthony, C., Goonetilleke, N., Karim, S.A., Margolis, D.M., Pond, S.K., Williamson, C., Swanstrom, R., 2019. The replication-competent HIV-1 latent reservoir is primarily established near the time of therapy initiation. Science Translational Medicine 11. https://doi.org/10.1126/scitranslmed.aaw5589

2. Archin, N.M., Kirchherr, J.L., Sung, J.A., Clutton, G., Sholtis, K., Xu, Y., Allard, B., Stuelke, E., Kashuba, A.D., Kuruc, J.D., Eron, J., Gay, C.L., Goonetilleke, N., Margolis, D.M., 2017. Interval dosing with the HDAC inhibitor vorinostat effectively reverses HIV latency. J. Clin. Invest. 127, 3126–3135. https://doi.org/10.1172/JCI92684

3. Arumugam, T., Ramphal, U., Adimulam, T., Chinniah, R., Ramsuran, V., 2021. Deciphering DNA Methylation in HIV Infection. Front Immunol 12, 795121. https://doi.org/10.3389/fimmu.2021.795121

4. Bailey, J.R., Sedaghat, A.R., Kieffer, T., Brennan, T., Lee, P.K., Wind-Rotolo, M., Haggerty, C.M., Kamireddi, A.R., Liu, Y., Lee, J., Persaud, D., Gallant, J.E., Cofrancesco, J., Quinn, T.C., Wilke, C.O., Ray, S.C., Siliciano, J.D., Nettles, R.E., Siliciano, R.F., 2006. Residual human immunodeficiency virus type 1 viremia in some patients on antiretroviral therapy is dominated by a small number of invariant clones rarely found in circulating CD4+ T cells. J Virol 80, 6441–6457. https://doi.org/10.1128/JVI.00591-06

5. Bartholomeeusen, K., Fujinaga, K., Xiang, Y., Peterlin, B.M., 2013. Histone Deacetylase Inhibitors (HDACis) That Release the Positive Transcription Elongation Factor b (P-TEFb) from Its Inhibitory Complex Also Activate HIV Transcription. Journal of Biological Chemistry 288, 14400–14407. https://doi.org/10.1074/jbc.M113.464834

6. Barton, K.M., Archin, N.M., Keedy, K.S., Espeseth, A.S., Zhang, Y., Gale, J., Wagner, F.F., Holson, E.B., Margolis, D.M., 2014. Selective HDAC Inhibition for the Disruption of Latent HIV-1 Infection. PLOS ONE 9, e102684. https://doi.org/10.1371/journal.pone.0102684

7. Battivelli, E., Dahabieh, M.S., Abdel-Mohsen, M., Svensson, J.P., Tojal Da Silva, I., Cohn, L.B., Gramatica, A., Deeks, S., Greene, W.C., Pillai, S.K., Verdin, E., 2018. Distinct chromatin functional states correlate with HIV latency reactivation in infected primary CD4+ T cells. eLife 7, e34655. https://doi.org/10.7554/eLife.34655

8. Boehm, D., Ott, M., 2017. Host Methyltransferases and Demethylases: Potential New Epigenetic Targets for HIV Cure Strategies and Beyond. AIDS Res Hum Retroviruses 33, S-8-S-22. https://doi.org/10.1089/aid.2017.0180

9. Bradley, T., Ferrari, G., Haynes, B.F., Margolis, D.M., Browne, E.P., 2018. Single-Cell Analysis of Quiescent HIV Infection Reveals Host Transcriptional Profiles that Regulate Proviral Latency. Cell Reports 25, 107–117.e3. https://doi.org/10.1016/j.celrep.2018.09.020

10. Brodin, J., Zanini, F., Thebo, L., Lanz, C., Bratt, G., Neher, R.A., Albert, J.,2016. Establishment and stability of the latent HIV-1 DNA reservoir. eLife 5, e18889. https://doi.org/10.7554/eLife.18889

11. Budhiraja, S., Famiglietti, M., Bosque, A., Planelles, V., Rice, A.P., 2013. Cyclin T1 and CDK9 T-loop phosphorylation are downregulated during establishment of HIV-1 latency in primary resting memory CD4+ T cells. J Virol 87, 1211–1220. https://doi.org/10.1128/JVI.02413-12

12. Buzon, M.J., Sun, H., Li, C., Shaw, A., Seiss, K., Ouyang, Z., Martin-Gayo, E., Leng, J., Henrich, T.J., Li, J.Z., Pereyra, F., Zurakowski, R., Walker, B.D., Rosenberg, E.S., Yu, X.G., Lichterfeld, M., 2014. HIV-1 persistence in CD4+ T cells with stem cell-like properties. Nat Med 20, 139–142. https://doi.org/10.1038/nm.3445

13. Chen, J.D., Evans, R.M., 1995. A transcriptional co-repressor that interacts with nuclear hormone receptors. Nature 377, 454–457. https://doi.org/10.1038/377454a0

14. Chen, L., Fischle, W., Verdin, E., Greene, W.C., 2001. Duration of Nuclear NF-κB Action Regulated by Reversible Acetylation. Science 293, 1653–1657. https://doi.org/10.1126/science.1062374

15. Chun, T.-W., Finzi, D., Margolick, J., Chadwick, K., Schwartz, D., Siliciano, R.F., 1995. In vivo fate of HIV-1-infected T cells: Quantitative analysis of the transition to stable latency. Nat Med 1, 1284–1290. https://doi.org/10.1038/nm1295-1284

16. Chun, T.W., Stuyver, L., Mizell, S.B., Ehler, L.A., Mican, J.A., Baseler, M., Lloyd, A.L., Nowak, M.A., Fauci, A.S., 1997. Presence of an inducible HIV-1 latent reservoir during highly active antiretroviral therapy. Proc Natl Acad Sci U S A 94, 13193–13197. https://doi.org/10.1073/pnas.94.24.13193

17. Clausen, D.J., Liu, J., Yu, W., Duffy, J.L., Chung, C.C., Myers, R.W., Klein, D.J., Fells, J., Holloway, K., Wu, J., Wu, G., Howell, B.J., Barnard, R.J.O., Kozlowski, J., 2020. Development of a selective HDAC inhibitor aimed at reactivating the HIV latent reservoir. Bioorganic & Medicinal Chemistry Letters 30, 127367. https://doi.org/10.1016/j.bmcl.2020.127367

18. Clutton, G., Xu, Y., Baldoni, P.L., Mollan, K.R., Kirchherr, J., Newhard, W., Cox, K., Kuruc, J.D., Kashuba, A., Barnard, R., Archin, N., Gay, C.L., Hudgens, M.G., Margolis, D.M., Goonetilleke, N., 2016. The differential short- and long-term effects of HIV-1 latency-reversing agents on T cell function. Sci Rep 6, 30749. https://doi.org/10.1038/srep30749

19. Cohn, L.B., Chomont, N., Deeks, S.G., 2020. The Biology of the HIV-1 Latent Reservoir and Implications for Cure Strategies. Cell Host & Microbe 27, 519–530. https://doi.org/10.1016/j.chom.2020.03.014

20. Conrad, R.J., Fozouni, P., Thomas, S., Sy, H., Zhang, Q., Zhou, M.-M., Ott, M., 2017. The Short Isoform of BRD4 Promotes HIV-1 Latency by Engaging Repressive SWI/SNF Chromatin Remodeling Complexes. Mol Cell 67, 1001–1012.e6. https://doi.org/10.1016/j.molcel.2017.07.025

21. Crooks, A.M., Bateson, R., Cope, A.B., Dahl, N.P., Griggs, M.K., Kuruc, J.D., Gay, C.L., Eron, J.J., Margolis, D.M., Bosch, R.J., Archin, N.M., 2015. Precise Quantitation of the Latent HIV-1 Reservoir: Implications for Eradication Strategies. The Journal of Infectious Diseases 212, 1361–1365. https://doi.org/10.1093/infdis/jiv218

22. Dobin, A., Davis, C.A., Schlesinger, F., Drenkow, J., Zaleski, C., Jha, S., Batut, P., Chaisson, M., Gingeras, T.R., 2013. STAR: ultrafast universal RNA-seq aligner. Bioinformatics 29, 15–21. https://doi.org/10.1093/bioinformatics/bts635

23. Doetzlhofer, A., Rotheneder, H., Lagger, G., Koranda, M., Kurtev, V., Brosch, G., Wintersberger, E., Seiser, C., 1999. Histone deacetylase 1 can repress transcription by binding to Sp1. Mol Cell Biol 19, 5504–5511. https://doi.org/10.1128/MCB.19.8.5504

24. Duette, G., Hiener, B., Morgan, H., Mazur, F.G., Mathivanan, V., Horsburgh, B.A., Fisher, K., Tong, O., Lee, E., Ahn, H., Shaik, A., Fromentin, R., Hoh, R., Bacchus-Souffan, C., Nasr, N., Cunningham, A., Hunt, P.W., Chomont, N., Turville, S.G., Deeks, S.G., Kelleher, A.D., Schlub, T.E., Palmer, S., 2022. The HIV-1 proviral landscape reveals Nef contributes to HIV-1 persistence in effector memory CD4+ T-cells. J Clin Invest. https://doi.org/10.1172/JCI154422

25. Einkauf, K.B., Osborn, M.R., Gao, C., Sun, W., Sun, X., Lian, X., Parsons, E.M., Gladkov, G.T., Seiger, K.W., Blackmer, J.E., Jiang, C., Yukl, S.A., Rosenberg, E.S., Yu, X.G., Lichterfeld, M., 2022. Parallel analysis of transcription, integration, and sequence of single HIV-1 proviruses. Cell 185, 266–282.e15. https://doi.org/10.1016/j.cell.2021.12.011

26. Elsheikh, M.M., Tang, Y., Li, D., Jiang, G., 2019. Deep latency: A new insight into a functional HIV cure. eBioMedicine 45, 624–629. https://doi.org/10.1016/j.ebiom.2019.06.020

27. Falcinelli, S.D., Kilpatrick, K.W., Read, J., Murtagh, R., Allard, B., Ghofrani, S., Kirchherr, J., James, K.S., Stuelke, E., Baker, C., Kuruc, J.D., Eron, J.J., Hudgens, M.G., Gay, C.L., Margolis, D.M., Archin, N.M., 2021. Longitudinal Dynamics of Intact HIV Proviral DNA and Outgrowth Virus Frequencies in a Cohort of Individuals Receiving Antiretroviral Therapy. J Infect Dis 224, 92–100. https://doi.org/10.1093/infdis/jiaa718

28. Fidler, S., Stöhr, W., Pace, Matt, Dorrell, L., Lever, A., Pett, S., Kinloch-de Loes, S., Fox, J., Clarke, A., Nelson, M., Thornhill, J., Khan, M., Fun, A., Bandara, M., Kelly, D., Kopycinski, J., Hanke, T., Yang, H., Bennett, R., Johnson, M., Howell, B., Barnard, R., Wu, G., Kaye, Steve, Wills, M., Babiker, A., Frater, J., Sandström, E., Darbyshire, J., Post, F., Conlon, C., Anderson, J., Maini, M., Peto, T., Sasieni, P., Miller, V., Weller, I., Fidler, S., Frater, J., Babiker, A., Stöhr, W., Pett, S., Dorrell, L., Pace, Matthew, Olejniczak, N., Brown, H., Robinson, N., Kopycinski, J., Yang, H., Hanke, T., Crook, A., Kaye, Steven, McClure, M., Erlwein, O., Lovell, A., Khan, M., Gabrielle, M., Bennett, R., Sy, A., Gregory, A., Hudson, F., Russell, C., Wood, G., Box, H., Kingsley, C., Topping, K., Lever, A., Wills, M., Fun, A., Bandara, M., Kelly, D., Collins, S., Markham, A., Rauchenberger, M., Sowunmi, Y., Shidfar, S., Hague, D., Nelson, M., Cerrone, M., Castrillo Martinez, N., Barber, T., Schoolmeesters, A., Weaver, C., Thunder, O., Rowlands, J., Higgs, C., Fedele, S., Bracchi, M., Thomas, L., Bourke, P., Nwokolo, N., Lawrenson, G., Fiorino, M., Lukha, H., Kinloch-de Loes, S., Johnson, M., Nightingale, A., Ngwu, N., Byrne, P., Cuthbertson, Z., Jones, M., Fernandez, T., Clarke, A., Fisher, M., Gleig, R., Trevitt, V., Fitzpatrick, C., Adams, T., Finnerty, F., Thornhill, J., Lewis, H., Kuldanek, K., Fox, J., Lwanga, J., Uzu, H., Lee, M., Merle, S., O’Rourke, P., Jendrulek, I., ZarkoFlynn, T., Taylor, M., Tiraboschi, J.M., Murray, T., 2020. Antiretroviral therapy alone versus antiretroviral therapy with a kick and kill approach, on measures of the HIV reservoir in participants with recent HIV infection (the RIVER trial): a phase 2, randomised trial. The Lancet 395, 888–898. https://doi.org/10.1016/S0140-6736(19)32990-3

29. Finzi, D., Blankson, J., Siliciano, J.D., Margolick, J.B., Chadwick, K., Pierson, T., Smith, K., Lisziewicz, J., Lori, F., Flexner, C., Quinn, T.C., Chaisson, R.E., Rosenberg, E., Walker, B., Gange, S., Gallant, J., Siliciano, R.F., 1999. Latent infection of CD4 + T cells provides a mechanism for lifelong persistence of HIV-1, even in patients on effective combination therapy. Nat Med 5, 512–517. https://doi.org/10.1038/8394

30. Gagne, M., Michaels, D., Lester, G.M.S., Gummuluru, S., Wong, W.W., Henderson, A.J., 2019. Strength of T cell signaling regulates HIV-1 replication and establishment of latency. PLOS Pathogens 15, e1007802. https://doi.org/10.1371/journal.ppat.1007802

31. Gay, C.L., James, K.S., Tuyishime, M., Falcinelli, S.D., Joseph, S.B., Moeser, M.J., Allard, B., Kirchherr, J.L., Clohosey, M., Raines, S.L.M., Montefiori, D.C., Shen, X., Gorelick, R.J., Gama, L., McDermott, A.B., Koup, R.A., Mascola, J.R., Floris-Moore, M., Kuruc, J.D., Ferrari, G., Eron, J.J., Archin, N.M., Margolis, D.M., 2022. Stable Latent HIV Infection and Low-level Viremia Despite Treatment With the Broadly Neutralizing Antibody VRC07-523LS and the Latency Reversal Agent Vorinostat. The Journal of Infectious Diseases 225, 856–861. https://doi.org/10.1093/infdis/jiab487

32. Gay, C.L., Kuruc, J.D., Falcinelli, S.D., Warren, J.A., Reifeis, S.A., Kirchherr, J.L., James, K.S., Dewey, M.G., Helms, A., Allard, B., Stuelke, E., Gamble, A., Plachco, A., Gorelick, R.J., Eron, J.J., Hudgens, M., Garrido, C., Goonetilleke, N., DeBenedette, M.A., Tcherepanova, I.Y., Nicolette, C.A., Archin, N.M., Margolis, D.M., 2020. Assessing the impact of AGS-004, a dendritic cell-based immunotherapy, and vorinostat on persistent HIV-1 Infection. Sci Rep 10, 5134. https://doi.org/10.1038/s41598-020-61878-3

33. Gillespie, M.A., Palii, C.G., Sanchez-Taltavull, D., Shannon, P., Longabaugh, W.J.R., Downes, D.J., Sivaraman, K., Espinoza, H.M., Hughes, J.R., Price, N.D., Perkins, T.J., Ranish, J.A., Brand, M. 2020. Absolute Quantification of Transcription Factors Reveals Principles of Gene Regulation in Erythropoiesis. Molecular Cell 78, 960–974.e11. https://doi.org/10.1016/j.molcel.2020.03.031

34. Goonetilleke, N., Clutton, G., Swanstrom, R., Joseph, S.B., 2019. Blocking Formation of the Stable HIV Reservoir: A New Perspective for HIV-1 Cure. Front Immunol 10, 1966. https://doi.org/10.3389/fimmu.2019.01966

35. Ho, Y.-C., Shan, L., Hosmane, N.N., Wang, J., Laskey, S.B., Rosenbloom, D.I.S., Lai, J., Blankson, J.N., Siliciano, J.D., Siliciano, R.F., 2013. Replication-Competent Noninduced Proviruses in the Latent Reservoir Increase Barrier to HIV-1 Cure. Cell 155, 540–551. https://doi.org/10.1016/j.cell.2013.09.020

36. Hsiao, F., Frouard, J., Gramatica, A., Xie, G., Telwatte, S., Lee, G.Q., Roychoudhury, P., Schwarzer, R., Luo, X., Yukl, S.A., Lee, S., Hoh, R., Deeks, S.G., Jones, R.B., Cavrois, M., Greene, W.C., Roan, N.R., 2020. Tissue memory CD4+ T cells expressing IL-7 receptor-alpha (CD127) preferentially support latent HIV-1 infection. PLOS Pathogens 16, e1008450. https://doi.org/10.1371/journal.ppat.1008450

37. Imai, K., Togami, H., Okamoto, T., 2010. Involvement of Histone H3 Lysine 9 (H3K9) Methyltransferase G9a in the Maintenance of HIV-1 Latency and Its Reactivation by BIX01294*. Journal of Biological Chemistry 285, 16538–16545. https://doi.org/10.1074/jbc.M110.103531

38. Jamaladdin, S., Kelly, R.D.W., O’Regan, L., Dovey, O.M., Hodson, G.E., Millard, C.J., Portolano, N., Fry, A.M., Schwabe, J.W.R., Cowley, S.M., 2014. Histone deacetylase (HDAC) 1 and 2 are essential for accurate cell division and the pluripotency of embryonic stem cells. PNAS 111, 9840–9845. https://doi.org/10.1073/pnas.1321330111

39. Jefferys, S.R., Burgos, S.D., Peterson, J.J., Selitsky, S.R., Turner, A.-M.W., James, L.I., Tsai, Y.-H., Coffey, A.R., Margolis, D.M., Parker, J., Browne, E.P., 2021. Epigenomic characterization of latent HIV infection identifies latency regulating transcription factors. PLOS Pathogens 17, e1009346. https://doi.org/10.1371/journal.ppat.1009346

40. Jiang, C., Lian, X., Gao, C., Sun, X., Einkauf, K.B., Chevalier, J.M., Chen, S.M.Y., Hua, S., Rhee, B., Chang, K., Blackmer, J.E., Osborn, M., Peluso, M.J., Hoh, R., Somsouk, M., Milush, J., Bertagnolli, L.N., Sweet, S.E., Varriale, J.A., Burbelo, P.D., Chun, T.-W., Laird, G.M., Serrao, E., Engelman, A.N., Carrington, M., Siliciano, R.F., Siliciano, J.M., Deeks, S.G., Walker, B.D., Lichterfeld, M., Yu, X.G., 2020. Distinct viral reservoirs in individuals with spontaneous control of HIV-1. Nature 585, 261–267. https://doi.org/10.1038/s41586-020-2651-8

41. Jones, B.R., Kinloch, N.N., Horacsek, J., Ganase, B., Harris, M., Harrigan, P.R., Jones, R.B., Brockman, M.A., Joy, J.B., Poon, A.F.Y., Brumme, Z.L., 2018. Phylogenetic approach to recover integration dates of latent HIV sequences within-host. Proc Natl Acad Sci U S A 115, E8958–E8967. https://doi.org/10.1073/pnas.1802028115

42. Kaya-Okur, H.S., Wu, S.J., Codomo, C.A., Pledger, E.S., Bryson, T.D., Henikoff, J.G., Ahmad, K., Henikoff, S., 2019. CUT&Tag for efficient epigenomic profiling of small samples and single cells. Nature Communications 10, 1930. https://doi.org/10.1038/s41467-019-09982-5

43. Keedy, K.S., Archin, N.M., Gates, A.T., Espeseth, A., Hazuda, D.J., Margolis, D.M., 2009. A limited group of class I histone deacetylases acts to repress human immunodeficiency virus type 1 expression. J. Virol. 83, 4749–4756. https://doi.org/10.1128/JVI.02585-08

44. Kroon, E.D.M.B., Ananworanich, J., Pagliuzza, A., Rhodes, A., Phanuphak, N., Trautmann, L., Mitchell, J.L., Chintanaphol, M., Intasan, J., Pinyakorn, S., Benjapornpong, K., Chang, J.J., Colby, D.J., Chomchey, N., Fletcher, J.L.K., Eubanks, K., Yang, H., Kapson, J., Dantanarayana, A., Tennakoon, S., Gorelick, R.J., Maldarelli, F., Robb, M.L., Kim, J.H., Spudich, S., Chomont, N., Phanuphak, P., Lewin, S.R., de Souza, M.S., SEARCH 019 and RV254 Study Teams, 2020. A randomized trial of vorinostat with treatment interruption after initiating antiretroviral therapy during acute HIV-1 infection. J Virus Erad 6, 100004. https://doi.org/10.1016/j.jve.2020.100004

45. Kulpa, D.A., Talla, A., Brehm, J.H., Ribeiro, S.P., Yuan, S., Bebin-Blackwell, A.-G., Miller, M., Barnard, R., Deeks, S.G., Hazuda, D., Chomont, N., Sékaly, R.-P., 2019. Differentiation into an Effector Memory Phenotype Potentiates HIV-1 Latency Reversal in CD4+ T Cells. Journal of Virology 93, e00969–19. https://doi.org/10.1128/JVI.00969-19

46. Kwon, K.J., Timmons, A.E., Sengupta, S., Simonetti, F.R., Zhang, H., Hoh, R., Deeks, S.G., Siliciano, J.D., Siliciano, R.F., 2020. Different human resting memory CD4+ T cell subsets show similar low inducibility of latent HIV-1 proviruses. Sci Transl Med 12, eaax6795. https://doi.org/10.1126/scitranslmed.aax6795

47. Lempicki, R.A., Kovacs, J.A., Baseler, M.W., Adelsberger, J.W., Dewar, R.L., Natarajan, V., Bosche, M.C., Metcalf, J.A., Stevens, R.A., Lambert, L.A., Alvord, W.G., Polis, M.A., Davey, R.T., Dimitrov, D.S., Lane, H.C., 2000. Impact of HIV-1 infection and highly active antiretroviral therapy on the kinetics of CD4+ and CD8+ T cell turnover in HIV-infected patients. Proc Natl Acad Sci U S A 97, 13778–13783.

48. Leus, N.G.J., Zwinderman, M.R.H., Dekker, F.J., 2016. Histone deacetylase 3 (HDAC 3) as emerging drug target in NF-κB-mediated inflammation. Curr Opin Chem Biol 33, 160– 168. https://doi.org/10.1016/j.cbpa.2016.06.019

49. Liberzon, A., Birger, C., Thorvaldsdóttir, H., Ghandi, M., Mesirov, J.P., Tamayo, P., 2015. The Molecular Signatures Database Hallmark Gene Set Collection. cels 1, 417–425. https://doi.org/10.1016/j.cels.2015.12.004

50. Liu, J., Yu, Y., Kelly, J., Sha, D., Alhassan, A.-B., Yu, W., Maletic, M.M., Duffy, J.L., Klein, D.J., Holloway, M.K., Carroll, S., Howell, B.J., Barnard, R.J.O., Wolkenberg, S., Kozlowski, J.A., 2020. Discovery of Highly Selective and Potent HDAC3 Inhibitors Based on a 2-Substituted Benzamide Zinc Binding Group. ACS Med Chem Lett 11, 2476–2483. https://doi.org/10.1021/acsmedchemlett.0c00462

51. Liu, R., Simonetti, F.R., Ho, Y.-C., 2020. The forces driving clonal expansion of the HIV-1 latent reservoir. Virol J 17, 4. https://doi.org/10.1186/s12985-019-1276-8

52. Lu, F., Zankharia, U., Vladimirova, O., Yi, Y., Collman, R.G., Lieberman, P.M., 2022. Epigenetic Landscape of HIV-1 Infection in Primary Human Macrophage. J Virol e0016222. https://doi.org/10.1128/jvi.00162-22

53. Lun, A.T.L., Smyth, G.K., 2016. csaw: a Bioconductor package for differential binding analysis of ChIP-seq data using sliding windows. Nucleic Acids Res 44, e45. https://doi.org/10.1093/nar/gkv1191

54. Mahnke, Y.D., Brodie, T.M., Sallusto, F., Roederer, M., Lugli, E., 2013. The who’s who of T-cell differentiation: human memory T-cell subsets. Eur. J. Immunol. 43, 2797–2809. https://doi.org/10.1002/eji.201343751

55. Maldarelli, F., Wu, X., Su, L., Simonetti, F.R., Shao, W., Hill, S., Spindler, J., Ferris, A.L., Mellors, J.W., Kearney, M.F., Coffin, J.M., Hughes, S.H., 2014. HIV latency. Specific HIV integration sites are linked to clonal expansion and persistence of infected cells. Science 345, 179–183. https://doi.org/10.1126/science.1254194

56. Malvaez, M., McQuown, S.C., Rogge, G.A., Astarabadi, M., Jacques, V., Carreiro, S., Rusche, J.R., Wood, M.A., 2013. HDAC3-selective inhibitor enhances extinction of cocaine-seeking behavior in a persistent manner. Proc Natl Acad Sci U S A 110, 2647–2652. https://doi.org/10.1073/pnas.1213364110

57. Manganaro, L., Hong, P., Hernandez, M.M., Argyle, D., Mulder, L.C.F., Potla, U., Diaz-Griffero, F., Lee, B., Fernandez-Sesma, A., Simon, V., 2018. IL-15 regulates susceptibility of CD4^+^ T cells to HIV infection. Proc Natl Acad Sci USA 115, E9659–E9667. https://doi.org/10.1073/pnas.1806695115

58. Margolis, D.M., 2011. Histone deacetylase inhibitors and HIV latency. Curr Opin HIV AIDS 6, 25–29. https://doi.org/10.1097/COH.0b013e328341242d

59. Margolis, D.M., Archin, N.M., Cohen, M.S., Eron, J.J., Ferrari, G., Garcia, J.V., Gay, C.L., Goonetilleke, N., Joseph, S.B., Swanstrom, R., Turner, A.-M.W., Wahl, A., 2020. Curing HIV: Seeking to Target and Clear Persistent Infection. Cell 181, 189–206. https://doi.org/10.1016/j.cell.2020.03.005

60. Martin, M., 2011. Cutadapt removes adapter sequences from high-throughput sequencing reads. EMBnet.journal 17, 10–12. https://doi.org/10.14806/ej.17.1.200

61. McManus, W.R., Bale, M.J., Spindler, J., Wiegand, A., Musick, A., Patro, S.C., Sobolewski, M.D., Musick, V.K., Anderson, E.M., Cyktor, J.C., Halvas, E.K., Shao, W., Wells, D., Wu, X., Keele, B.F., Milush, J.M., Hoh, R., Mellors, J.W., Hughes, S.H., Deeks, S.G., Coffin, J.M., Kearney, M.F., 2019. HIV-1 in lymph nodes is maintained by cellular proliferation during antiretroviral therapy. J Clin Invest 129, 4629–4642. https://doi.org/10.1172/JCI126714

62. Mediouni, S., Chinthalapudi, K., Ekka, M.K., Usui, I., Jablonski, J.A., Clementz, M.A., Mousseau, G., Nowak, J., Macherla, V.R., Beverage, J.N., Esquenazi, E., Baran, P., de Vera, I.M.S., Kojetin, D., Loret, E.P., Nettles, K., Maiti, S., Izard, T., Valente, S.T., 2019. Didehydro-Cortistatin A Inhibits HIV-1 by Specifically Binding to the Unstructured Basic Region of Tat. mBio. https://doi.org/10.1128/mBio.02662-18

63. Meers, M.P., Bryson, T.D., Henikoff, J.G., Henikoff, S., 2019. Improved CUT&RUN chromatin profiling tools. eLife 8, e46314. https://doi.org/10.7554/eLife.46314

64. Mohri, H., Perelson, A.S., Tung, K., Ribeiro, R.M., Ramratnam, B., Markowitz, M., Kost, R., Hurley Weinberger, L., Cesar, D., Hellerstein, M.K., Ho, D.D., 2001. Increased Turnover of T Lymphocytes in HIV-1 Infection and Its Reduction by Antiretroviral Therapy. J Exp Med 194, 1277–1288.

65. Murray, A.J., Kwon, K.J., Farber, D.L., Siliciano, R.F., 2016. The Latent Reservoir for HIV-1: How Immunologic Memory and Clonal Expansion Contribute to HIV-1 Persistence. J.I. 197, 407–417. https://doi.org/10.4049/jimmunol.1600343

66. Nabel, G., Baltimore, D., 1987. An inducible transcription factor activates expression of human immunodeficiency virus in T cells. Nature 326, 711–713. https://doi.org/10.1038/326711a0

67. Narita, Masashi, Nuñez, S., Heard, E., Narita, Masako, Lin, A.W., Hearn, S.A., Spector, D.L., Hannon, G.J., Lowe, S.W., 2003. Rb-Mediated Heterochromatin Formation and Silencing of E2F Target Genes during Cellular Senescence. Cell 113, 703–716. https://doi.org/10.1016/S0092-8674(03)00401-X

68. Nguyen, D.N., Roth, T.L., Li, P.J., Chen, P.A., Apathy, R., Mamedov, M.R., Vo, L.T., Tobin, V.R., Goodman, D., Shifrut, E., Bluestone, J.A., Puck, J.M., Szoka, F.C., Marson, A., 2020. Polymer-stabilized Cas9 nanoparticles and modified repair templates increase genome editing efficiency. Nature Biotechnology 38, 44–49. https://doi.org/10.1038/s41587-019-0325-6

69. Nguyen, H.C.B., Adlanmerini, M., Hauck, A.K., Lazar, M.A., 2020. Dichotomous engagement of HDAC3 activity governs inflammatory responses. Nature 584, 286–290. https://doi.org/10.1038/s41586-020-2576-2

70. Nixon, C.C., Mavigner, M., Sampey, G.C., Brooks, A.D., Spagnuolo, R.A., Irlbeck, D.M., Mattingly, C., Ho, P.T., Schoof, N., Cammon, C.G., Tharp, G.K., Kanke, M., Wang, Z., Cleary, R.A., Upadhyay, A.A., De, C., Wills, S.R., Falcinelli, S.D., Galardi, C., Walum, H., Schramm, N.J., Deutsch, J., Lifson, J.D., Fennessey, C.M., Keele, B.F., Jean, S., Maguire, S., Liao, B., Browne, E.P., Ferris, R.G., Brehm, J.H., Favre, D., Vanderford, T.H., Bosinger, S.E., Jones, C.D., Routy, J.-P., Archin, N.M., Margolis, D.M., Wahl, A., Dunham, R.M., Silvestri, G., Chahroudi, A., Garcia, J.V., 2020. Systemic HIV and SIV latency reversal via non-canonical NF-κB signalling in vivo. Nature 578, 160–165. https://doi.org/10.1038/s41586-020-1951-3

71. Oh, S.A., Seki, A., Rutz, S., 2019. Ribonucleoprotein Transfection for CRISPR/Cas9-Mediated Gene Knockout in Primary T Cells. Curr Protoc Immunol 124, e69. https://doi.org/10.1002/cpim.69

72. Ophinni, Y., Inoue, M., Kotaki, T., Kameoka, M., 2018. CRISPR/Cas9 system targeting regulatory genes of HIV-1 inhibits viral replication in infected T-cell cultures. Scientific Reports 8, 7784. https://doi.org/10.1038/s41598-018-26190-1

73. Park, S.-Y., Kim, J.-S., 2020. A short guide to histone deacetylases including recent progress on class II enzymes. Exp Mol Med 52, 204–212. https://doi.org/10.1038/s12276-020-0382-4

74. Pearson, R., Kim, Y.K., Hokello, J., Lassen, K., Friedman, J., Tyagi, M., Karn, J., 2008. Epigenetic Silencing of Human Immunodeficiency Virus (HIV) Transcription by Formation of Restrictive Chromatin Structures at the Viral Long Terminal Repeat Drives the Progressive Entry of HIV into Latency. J Virol 82, 12291–12303. https://doi.org/10.1128/JVI.01383-08

75. Perelson, A.S., Essunger, P., Cao, Y., Vesanen, M., Hurley, A., Saksela, K., Markowitz, M., Ho, D.D., 1997. Decay characteristics of HIV-1-infected compartments during combination therapy. Nature 387, 188–191. https://doi.org/10.1038/387188a0

76. Pinzone, M.R., VanBelzen, D.J., Weissman, S., Bertuccio, M.P., Cannon, L., Venanzi-Rullo, E., Migueles, S., Jones, R.B., Mota, T., Joseph, S.B., Groen, K., Pasternak, A.O., Hwang, W.-T., Sherman, B., Vourekas, A., Nunnari, G., O’Doherty, U., 2019. Longitudinal HIV sequencing reveals reservoir expression leading to decay which is obscured by clonal expansion. Nat Commun 10. https://doi.org/10.1038/s41467-019-08431-7

77. Quinlan, A.R., Hall, I.M., 2010. BEDTools: a flexible suite of utilities for comparing genomic features. Bioinformatics 26, 841–842. https://doi.org/10.1093/bioinformatics/btq033

78. Rafati, H., Parra, M., Hakre, S., Moshkin, Y., Verdin, E., Mahmoudi, T., 2011. Repressive LTR Nucleosome Positioning by the BAF Complex Is Required for HIV Latency. PLOS Biology 9, e1001206. https://doi.org/10.1371/journal.pbio.1001206

79. Ramalingam, S.S., Parise, R.A., Ramananthan, R.K., Lagattuta, T.F., Musguire, L.A., Stoller, R.G., Potter, D.M., Argiris, A.E., Zwiebel, J.A., Egorin, M.J., Belani, C.P., 2007. Phase I and Pharmacokinetic Study of Vorinostat, A Histone Deacetylase Inhibitor, in Combination with Carboplatin and Paclitaxel for Advanced Solid Malignancies. Clinical Cancer Research 13, 3605–3610. https://doi.org/10.1158/1078-0432.CCR-07-0162

80. Ramírez, F., Ryan, D.P., Grüning, B., Bhardwaj, V., Kilpert, F., Richter, A.S., Heyne, S., Dündar, F., Manke, T., 2016. deepTools2: a next generation web server for deep-sequencing data analysis. Nucleic Acids Res 44, W160–165. https://doi.org/10.1093/nar/gkw257

81. Rasmussen, T.A., Tolstrup, M., Brinkmann, C.R., Olesen, R., Erikstrup, C., Solomon, A., Winckelmann, A., Palmer, S., Dinarello, C., Buzon, M., Lichterfeld, M., Lewin, S.R., Østergaard, L., Søgaard, O.S., 2014. Panobinostat, a histone deacetylase inhibitor, for latent-virus reactivation in HIV-infected patients on suppressive antiretroviral therapy: a phase 1/2, single group, clinical trial. The Lancet HIV 1, e13–e21. https://doi.org/10.1016/S2352-3018(14)70014-1

82. Romerio, F., Gabriel, M.N., Margolis, D.M., 1997. Repression of human immunodeficiency virus type 1 through the novel cooperation of human factors YY1 and LSF. J Virol 71, 9375– 9382.

83. Roux, A., Leroy, H., Muylder, B.D., Bracq, L., Oussous, S., Dusanter-Fourt, I., Chougui, G., Tacine, R., Randriamampita, C., Desjardins, D., Grand, R.L., Bouillaud, F., Benichou, S., Margottin-Goguet, F., Cheynier, R., Bismuth, G., Mangeney, M., 2019. FOXO1 transcription factor plays a key role in T cell—HIV-1 interaction. PLOS Pathogens 15, e1007669. https://doi.org/10.1371/journal.ppat.1007669

84. Sahu, G.K., Lee, K., Ji, J., Braciale, V., Baron, S., Cloyd, M.W., 2006. A novel in vitro system to generate and study latently HIV-infected long-lived normal CD4+ T-lymphocytes. Virology 355, 127–137. https://doi.org/10.1016/j.virol.2006.07.020

85. Seki, A., Rutz, S., 2018. Optimized RNP transfection for highly efficient CRISPR/Cas9-mediated gene knockout in primary T cells. J. Exp. Med. 215, 985–997. https://doi.org/10.1084/jem.20171626

86. Shan, L., Deng, K., Gao, H., Xing, S., Capoferri, A.A., Durand, C.M., Rabi, S.A., Laird, G.M., Kim, M., Hosmane, N.N., Yang, H.-C., Zhang, H., Margolick, J.B., Li, L., Cai, W., Ke, R., Flavell, R.A., Siliciano, J.D., Siliciano, R.F., 2017. Transcriptional Reprogramming during Effector-to-Memory Transition Renders CD4+ T Cells Permissive for Latent HIV-1 Infection. Immunity 47, 766–775.e3. https://doi.org/10.1016/j.immuni.2017.09.014

87. Shan, L., Xing, S., Yang, H.-C., Zhang, H., Margolick, J.B., Siliciano, R.F., 2014. Unique characteristics of histone deacetylase inhibitors in reactivation of latent HIV-1 in Bcl-2-transduced primary resting CD4+ T cells. J Antimicrob Chemother 69, 28–33. https://doi.org/10.1093/jac/dkt338

88. Siliciano, J.D., Kajdas, J., Finzi, D., Quinn, T.C., Chadwick, K., Margolick, J.B., Kovacs, C., Gange, S.J., Siliciano, R.F., 2003. Long-term follow-up studies confirm the stability of the latent reservoir for HIV-1 in resting CD4+ T cells. Nat Med 9, 727–728. https://doi.org/10.1038/nm880

89. Siliciano, J.D., Siliciano, R.F., 2021. Low Inducibility of Latent Human Immunodeficiency Virus Type 1 Proviruses as a Major Barrier to Cure. J Infect Dis 223, 13–21. https://doi.org/10.1093/infdis/jiaa649

90. Simonetti, F.R., Zhang, H., Soroosh, G.P., Duan, J., Rhodehouse, K., Hill, A.L., Beg, S.A., McCormick, K., Raymond, H.E., Nobles, C.L., Everett, J.K., Kwon, K.J., White, J.A., Lai, J., Margolick, J.B., Hoh, R., Deeks, S.G., Bushman, F.D., Siliciano, J.D., Siliciano, R.F., 2021. Antigen-driven clonal selection shapes the persistence of HIV-1–infected CD4^+^ T cells in vivo. J Clin Invest 131. https://doi.org/10.1172/JCI145254

91. Skene, P.J., Henikoff, S., 2017. An efficient targeted nuclease strategy for high-resolution mapping of DNA binding sites. eLife 6, e21856. https://doi.org/10.7554/eLife.21856

92. Soriano-Sarabia, N., Bateson, R.E., Dahl, N.P., Crooks, A.M., Kuruc, J.D., Margolis, D.M., Archin, N.M., 2014. Quantitation of replication-competent HIV-1 in populations of resting CD4+ T cells. J. Virol. 88, 14070–14077. https://doi.org/10.1128/JVI.01900-14

93. Spina, C.A., Anderson, J., Archin, N.M., Bosque, A., Chan, J., Famiglietti, M., Greene, W.C., Kashuba, A., Lewin, S.R., Margolis, D.M., Mau, M., Ruelas, D., Saleh, S., Shirakawa, K., Siliciano, R.F., Singhania, A., Soto, P.C., Terry, V.H., Verdin, E., Woelk, C., Wooden, S., Xing, S., Planelles, V., 2013. An In-Depth Comparison of Latent HIV-1 Reactivation in Multiple Cell Model Systems and Resting CD4+ T Cells from Aviremic Patients. PLOS Pathogens 9, e1003834. https://doi.org/10.1371/journal.ppat.1003834

94. Spina, C.A., Guatelli, J.C., Richman, D.D., 1995. Establishment of a stable, inducible form of human immunodeficiency virus type 1 DNA in quiescent CD4 lymphocytes in vitro. J Virol 69, 2977–2988. https://doi.org/10.1128/JVI.69.5.2977-2988.1995

95. Spina, C.A., Prince, H.E., Richman, D.D., 1997. Preferential replication of HIV-1 in the CD45RO memory cell subset of primary CD4 lymphocytes in vitro. J Clin Invest 99, 1774–1785. https://doi.org/10.1172/JCI119342

96. Sun, Z., Feng, D., Fang, B., Mullican, S.E., You, S.-H., Lim, H.-W., Everett, L.J., Nabel, C.S., Li, Y., Selvakumaran, V., Won, K.-J., Lazar, M.A., 2013. Deacetylase-Independent Function of HDAC3 in Transcription and Metabolism Requires Nuclear Receptor Corepressor. Molecular Cell 52, 769–782. https://doi.org/10.1016/j.molcel.2013.10.022

97. Swiggard, W.J., Baytop, C., Yu, J.J., Dai, J., Li, C., Schretzenmair, R., Theodosopoulos, T., O’Doherty, U., 2005. Human Immunodeficiency Virus Type 1 Can Establish Latent Infection in Resting CD4+ T Cells in the Absence of Activating Stimuli. Journal of Virology 79, 14179–14188. https://doi.org/10.1128/JVI.79.22.14179-14188.2005

98. Ting, P.Y., Parker, A.E., Lee, J.S., Trussell, C., Sharif, O., Luna, F., Federe, G., Barnes, S.W., Walker, J.R., Vance, J., Gao, M.-Y., Klock, H.E., Clarkson, S., Russ, C., Miraglia, L.J., Cooke, M.P., Boitano, A.E., McNamara, P., Lamb, J., Schmedt, C., Snead, J.L., 2018. Guide Swap enables genome-scale pooled CRISPR–Cas9 screening in human primary cells. Nature Methods 15, 941–946. https://doi.org/10.1038/s41592-018-0149-1

99. Tripathy, M.K., McManamy, M.E.M., Burch, B.D., Archin, N.M., Margolis, D.M., 2015. H3K27 Demethylation at the Proviral Promoter Sensitizes Latent HIV to the Effects of Vorinostat in Ex Vivo Cultures of Resting CD4+ T Cells. Journal of Virology 89, 8392–8405. https://doi.org/10.1128/JVI.00572-15

100. Vallejo-Gracia, A., Chen, I.P., Perrone, R., Besnard, E., Boehm, D., Battivelli, E., Tezil, T., Krey, K., Raymond, K.A., Hull, P.A., Walter, M., Habrylo, I., Cruz, A., Deeks, S., Pillai, S., Verdin, E., Ott, M., 2020. FOXO1 promotes HIV latency by suppressing ER stress in T cells. Nat Microbiol. https://doi.org/10.1038/s41564-020-0742-9

101. van der Vlag, J., Otte, A.P., 1999. Transcriptional repression mediated by the human polycomb-group protein EED involves histone deacetylation. Nat Genet 23, 474–478. https://doi.org/10.1038/70602

102. Wagner, T.A., McKernan, J.L., Tobin, N.H., Tapia, K.A., Mullins, J.I., Frenkel, L.M., 2013. An increasing proportion of monotypic HIV-1 DNA sequences during antiretroviral treatment suggests proliferation of HIV-infected cells. J Virol 87, 1770–1778. https://doi.org/10.1128/JVI.01985-12

103. White, J.A., Simonetti, F.R., Beg, S., McMyn, N.F., Dai, W., Bachmann, N., Lai, J., Ford, W.C., Bunch, C., Jones, J.L., Ribeiro, R.M., Perelson, A.S., Siliciano, J.D., Siliciano, R.F., 2022. Complex decay dynamics of HIV virions, intact and defective proviruses, and 2LTR circles following initiation of antiretroviral therapy. Proc Natl Acad Sci U S A 119, e2120326119. https://doi.org/10.1073/pnas.2120326119

104. Williams, S.A., Chen, L.-F., Kwon, H., Ruiz-Jarabo, C.M., Verdin, E., Greene, W.C., 2006. NF-kappaB p50 promotes HIV latency through HDAC recruitment and repression of transcriptional initiation. EMBO J 25, 139–149. https://doi.org/10.1038/sj.emboj.7600900

105. Wilting, R.H., Yanover, E., Heideman, M.R., Jacobs, H., Horner, J., van der Torre, J., DePinho, R.A., Dannenberg, J.-H., 2010. Overlapping functions of Hdac1 and Hdac2 in cell cycle regulation and haematopoiesis. EMBO J 29, 2586–2597. https://doi.org/10.1038/emboj.2010.136

106. Wonderlich, E.R., Subramanian, K., Cox, B., Wiegand, A., Lackman-Smith, C., Bale, M.J., Stone, M., Hoh, R., Kearney, M.F., Maldarelli, F., Deeks, S.G., Busch, M.P., Ptak, R.G., Kulpa, D.A., 2019. Effector memory differentiation increases detection of replication-competent HIV-l in resting CD4+ T cells from virally suppressed individuals. PLOS Pathogens 15, e1008074. https://doi.org/10.1371/journal.ppat.1008074

107. Wu, S.J., Furlan, S.N., Mihalas, A.B., Kaya-Okur, H.S., Feroze, A.H., Emerson, S.N., Zheng, Y., Carson, K., Cimino, P.J., Keene, C.D., Sarthy, J.F., Gottardo, R., Ahmad, K., Henikoff, S., Patel, A.P., 2021. Single-cell CUT&Tag analysis of chromatin modifications in differentiation and tumor progression. Nat Biotechnol 39, 819–824. https://doi.org/10.1038/s41587-021-00865-z

108. Yang, H.-C., Xing, S., Shan, L., O’Connell, K., Dinoso, J., Shen, A., Zhou, Y., Shrum, C.K., Han, Y., Liu, J.O., Zhang, H., Margolick, J.B., Siliciano, R.F., 2009. Small-molecule screening using a human primary cell model of HIV latency identifies compounds that reverse latency without cellular activation. J Clin Invest 119, 3473–3486. https://doi.org/10.1172/JCI39199

109. Yu, W., Fells, J., Clausen, D., Liu, J., Klein, D.J., Christine Chung, C., Myers, R.W., Wu, J., Wu, G., Howell, B.J., Barnard, R.J.O., Kozlowski, J., 2021. Discovery of macrocyclic HDACs 1, 2, and 3 selective inhibitors for HIV latency reactivation. Bioorganic & Medicinal Chemistry Letters 47, 128168. https://doi.org/10.1016/j.bmcl.2021.128168

110. Yukawa, M., Jagannathan, S., Vallabh, S., Kartashov, A.V., Chen, X., Weirauch, M.T., Barski, A., 2019. AP-1 activity induced by co-stimulation is required for chromatin opening during T cell activation. Journal of Experimental Medicine 217, e20182009. https://doi.org/10.1084/jem.20182009

111. Zack, J.A., Arrigo, S.J., Weitsman, S.R., Go, A.S., Haislip, A., Chen, I.S., 1990. HIV-1 entry into quiescent primary lymphocytes: molecular analysis reveals a labile, latent viral structure. Cell 61, 213–222. https://doi.org/10.1016/0092-8674(90)90802-l

112. Zhang, Y., Liu, T., Meyer, C.A., Eeckhoute, J., Johnson, D.S., Bernstein, B.E., Nusbaum, C., Myers, R.M., Brown, M., Li, W., Liu, X.S., 2008. Model-based analysis of ChIP-Seq (MACS). Genome Biol 9, R137. https://doi.org/10.1186/gb-2008-9-9-r137

